# Sex separation unveils the functional plasticity of the vomeronasal organ in rabbits

**DOI:** 10.1101/2022.08.24.505087

**Authors:** PR Villamayor, J Gullón, L Quintela, P Sánchez-Quinteiro, P Martínez, D Robledo

**Author notes:** Both authors contributed equally to this manuscript (P Martínez, D Robledo).

## Abstract

**Background:** Chemosensory cues are vital for social and sexual behaviours and are primarily detected and processed by the vomeronasal system (VNS), whose plastic capacity has been investigated in mice. However, studying chemosensory plasticity outside of laboratory conditions may give a more realistic picture of how the VNS adapts to a changing environment. Rabbits are a well-described model of chemocommunication since the discovery of the rabbit mammary pheromone and their vomeronasal organ (VNO) transcriptome was recently characterized, a first step to further study plasticity-mediated transcriptional changes. In this study, we assess the plastic capacity of the rabbit male and female VNO under sex-separation *vs* sex-combined scenarios, including adults and juveniles, to determine whether the rabbit VNO is plastic and, if so, whether such plasticity is already established at early stages of life.

**Results:** First, we characterized the number of differentially expressed genes (DEGs) between the VNO of rabbit male and female under sex-separation and compared it to sex-combined individuals, both in adults and juveniles, finding that differences between male and female were larger in a sex-separated scenario. Secondly, we analyzed the number of DEGs between sex-separated and sex-combined scenarios, both in males and females. In adults, both sexes showed a high number of DEGs while in juveniles only females showed differences. Additionally, the vomeronasal receptor genes were strikingly down-regulated in sex-separated adult females, whereas in juveniles up-regulation was shown for the same condition, suggesting a role of VRs in puberty onset. Finally, we described the environment-modulated plastic capacity of genes involved in reproduction, immunity and VNO functional activity, including G-protein coupled receptors.

**Conclusions:** Our results show that sex-separation induces sex- and stage- specific gene expression differences in the VNO of male and female rabbit, both in adults and juveniles. These results bring out for the first time the plastic capacity of the rabbit VNO, supporting its functional adaptation to specifically respond to a continuous changing environment. Finally, species-specific differences and individual variability should always be considered in VNO studies and overall chemocommunication research.

## Introduction

Chemosensory communication controls many vital social and sexual behaviors such as fighting or mating and strongly impacts individual behaviour, demanding accuracy and dynamic range of stimuli perception as well as their underlying transduction mechanisms. In mammals, chemical stimuli are predominantly detected through the VNS, considered as a hardwired system in charge of detecting innate and stereotyped chemosignals [1]. However, since a few decades ago, investigations have been pointing towards a more dynamic VNS system. For instance, its high degree of plasticity was described by Ichikawa [2], especially regarding synaptic morphology variation in the accessory olfactory bulb, the first center for vomeronasal signal integration within the central nervous system. Lately, the VNS has entered a ‘new era’ in which it was finally recognized as a system with capacity for adaptative learning, and experience- and state- dependent plasticity [3–7].

Chemical stimuli are released by one individual to the external world and detected by conspecifics, predominantly by vomeronasal receptors (VRs) located in vomeronasal sensory neurons (VSNs) within the vomeronasal organ (VNO) [8]. Vomeronasal receptors are highly fine-tuned to be able to respond efficiently to continuous environmental changes. The two main types of vomeronasal receptors, V1R and V2R, have been involved in the detection of physiological status of conspecifics by binding distinct classes of sexual steroids (V1R), and in sex-related behaviours (V2R), such as puberty acceleration, female attraction or male aggressiveness [9]. Additionally, in mice, a multigene family of non-classical class I major histocompatibility complex (MHC) genes (H2-Mv), were found co-expressed with V2R [10] and, even though their functionality has not been fully explored, they contribute to ultrasensitive chemodetection by a subset of VSNs [11]. Another type of vomeronasal receptors, representing an expansion of the family formyl peptide receptors (FPRs), has been identified in vomeronasal neurons in mice [12, 13]. This family is also expressed in immune cells directly involved in innate immunity and in recognition of several inflammation-related molecules [14–17]. Therefore, the vomeronasal system appears to be closely related to the immune system and its role in pathogen sensing [18], detection and avoidance of sick conspecifics [19] and genetic compatibility [20], has been previously established. Due to the wide diversity of species-specific chemical stimuli perceived by the VNO, it is likely that other unknown types of vomeronasal receptors might be found in the VNO. For instance, it has been reported that VSNs are highly sensitive to excreted sex-steroids, glucocorticoids and bile acids [21] although their plasma membrane receptors remain unclear. VNO cytoplasmatic sex-steroid receptors may arise as potential candidates to perceive the physiological state of a given individual.

The VNO also shows a high plasticity range to cope with environmental changes. Marom et al. [5] have recently determined that responses triggered by VSNs are highly plastic and greatly vary upon experience. Also, it is well-established that VSNs undergo regeneration throughout life both physiologically and after injury, thus supporting their plastic capacity [22, 23]. Importantly, experience-dependent plasticity triggers dramatic changes at the individual level in the abundance of specific functional types of VSNs, inducing specific behaviours [24]. Such individual variability adds further complexity to the study of the VNO. Experience has been documented to affect gene expression in VSNs, and, in particular, to that of vomeronasal receptors (VRs), but its correlation to neuron abundance has not been determined [25, 26]. Recently, long-term (6 months) exposure to a particular social environment (sex-separation conditions) has proved to induce changes in the gene expression of mice VNO both in the abundance (in situ hybridization) and gene expression (RNA sequencing, RNAseq) of VSNs and VRs [27, 28]. However, it remains unclear whether shorter exposure to this particular condition would follow a similar plasticity. Also, since gene expression points to the function and pathways of specific genes, we argue that transcriptional changes, regardless neuron abundance, might be the main mechanism by which olfactory sensory systems adapt to a particular environmental change. Indeed, despite only few studies have approached transcriptional changes in the VNO, data from the main olfactory epithelium strongly supports transcriptional changes underlying sensory adaptation [29]. Recently, it has been determined that olfactory sensory neurons adapt to a particular environment via the activation of highly specific transcriptomic programs [30]. Since the VNO is another sensorial structure which needs to adapt and react to particular scenarios, it seems reasonable that the VNO follows similar mechanisms of transcriptional adaptation.

Mice have been the gold standard species in chemocommunication studies. However, the species-specific nature of the VNS makes it necessary to broaden the species range to achieve reliable conclusions. Rabbits are considered a suitable model of chemocommunication since the discovery of the rabbit mammary pheromone [31]. A previous VNO transcriptomic study on this species demonstrated its complex nature, and expression of a wide range of VRs and other genes involved in communication and reproduction was reported [32]. In this study, we targeted for the first time the potential plasticity of the rabbit VNO. We used RNAseq to study the gene expression of the rabbit VNO under sex-separation and sex-combined conditions in adults (6 months old), considered as a long-term exposure and in which neural turnover is expected, and juveniles (40 days old), to decipher whether sensory plasticity starts at earlier stages. We found VNO plastic changes underlying sex-separation in male and female adults. Surprisingly, in juveniles, VNO plasticity was found in females but not in males, a feature that points to a potential relationship between the VNO activity and the onset of puberty. Special attention was focused on the VR repertoire, where we found striking down-regulation of VR genes in adult females when sex-separated. Other features such as expression changes of sex-steroid receptors or specific G protein coupled receptors were also detected. All in all, transcriptional modulation appears to play an important role in sensory adaptation, suggesting that the VNO may evolve to respond to the environment via transcription modulation since early life stages.

## Results

The main goal of this study was to analyze the plasticity and capacity for sensorial adaptation of the VNO in response to diverse socio-environmental conditions. To this end, male and female rabbits were separated at birth (sex-separated), and their VNO transcriptome in juveniles (40 days) and adults (6 months) was compared to that of males and females reared together since birth (sex-combined) (Fig. 1a; Additional Fig. 1).

**Fig 1.**
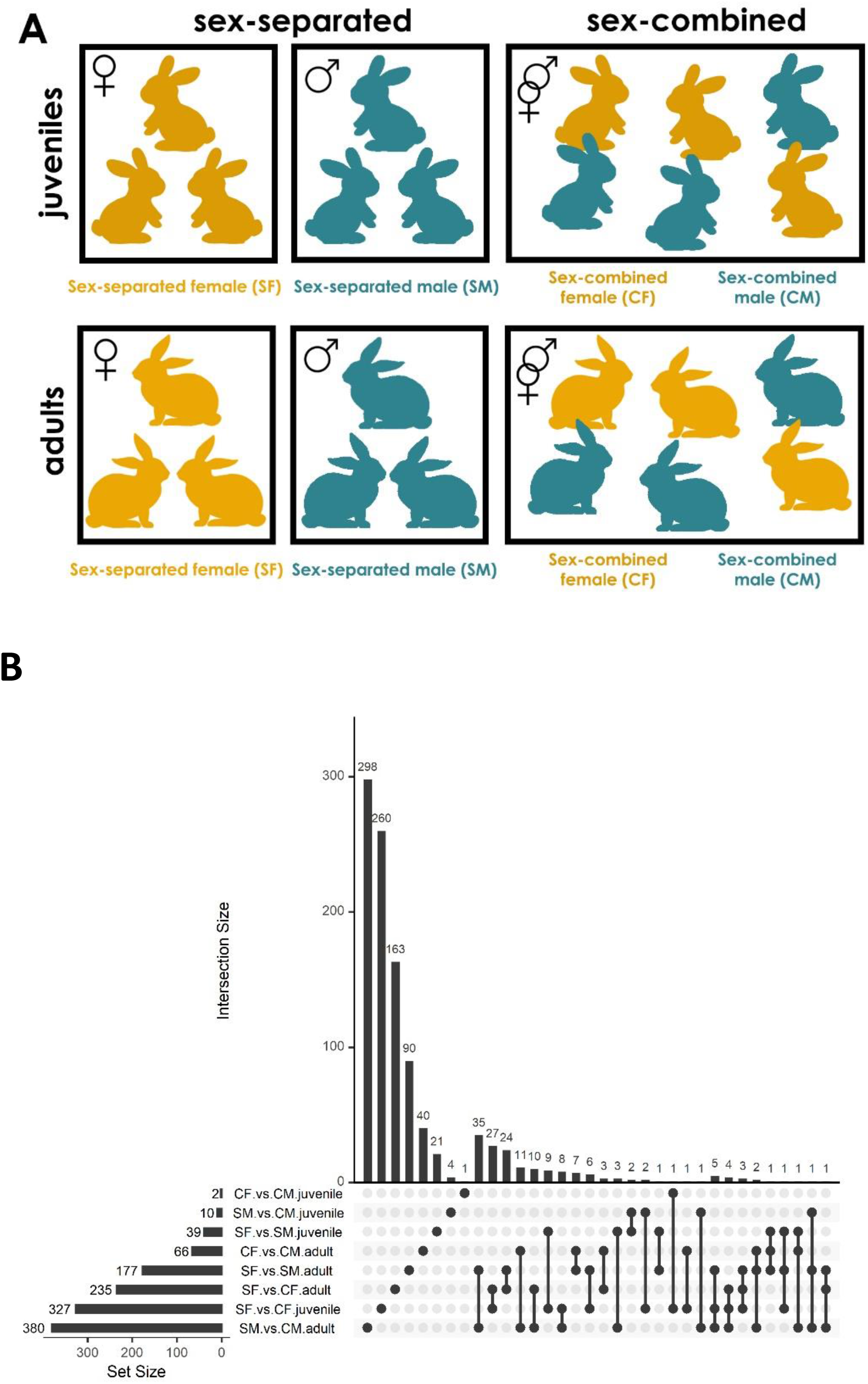
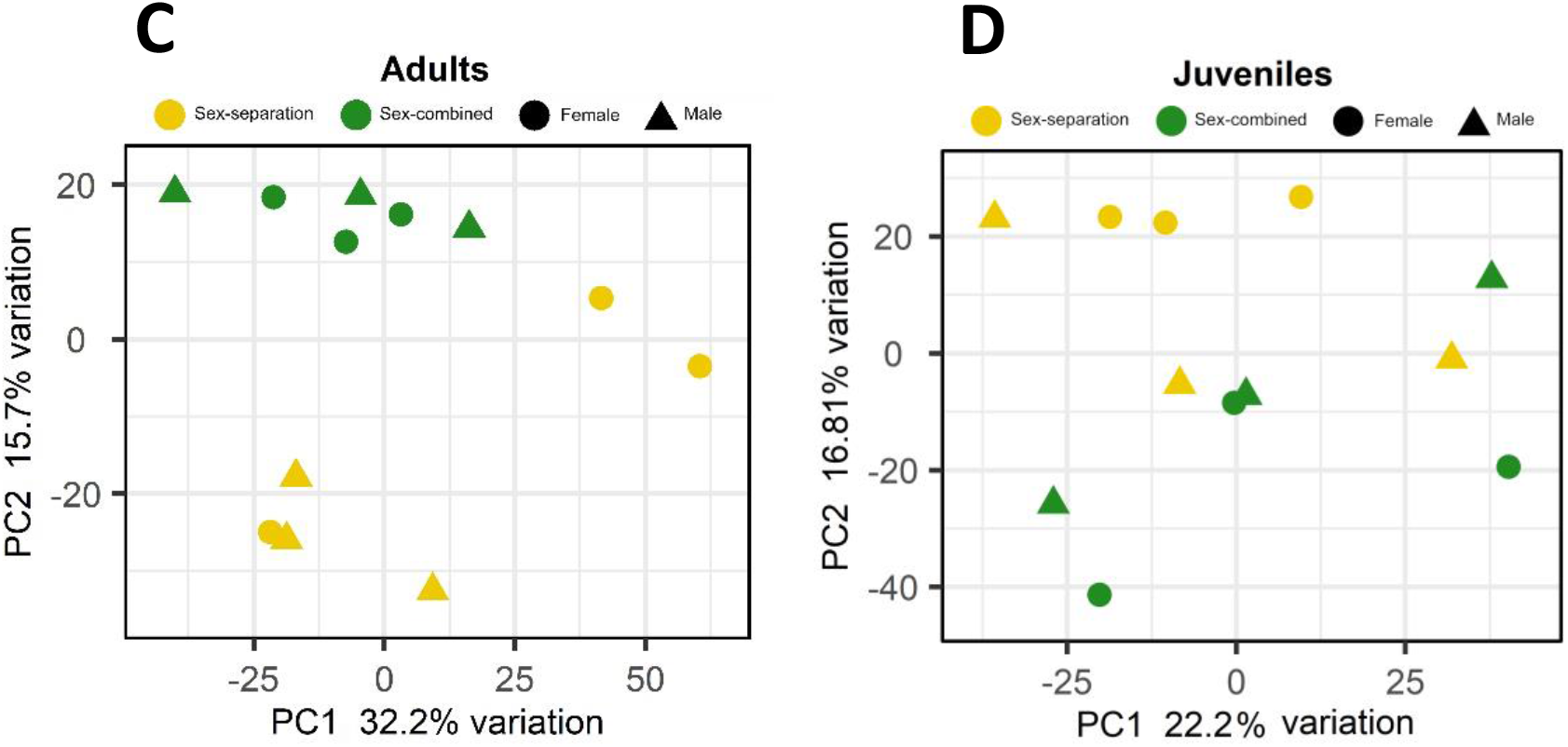
Gene expression differences between male and female rabbits in VNO tissue, both in adults and juveniles, under different socio-environmental conditions. **(a)** Experimental design. From day 0 until day 40 -first sampling point (juveniles)- and day 180 -second sampling point (adults)-, rabbits experienced either a sex-separated environment or a sex-combined environment. **(b)** Upsetplot showing the number of DEGs of each experimental group and those shared between groups. The bars representing the intersection size show the number of DEGs shared by the experimental groups highlighted by black dots in the matrix panel below. **(c, d)** Principal component analysis (PCA) where sex-separated and sex-combined males and females are plotted, both in (c) adults and (d) juveniles.

All animals were reared under the same farm environment and management protocols. On day 0, kits were re-distributed among mothers, so that each mother took care of 8 kits coming from different mothers. For the sex-separated condition, female and male kits were separated since birth in different litters. We employed 50 litters/condition (50 for sex-separated females and 50 for sex-separated males) to ensure that animals were surrounded by individuals under the same condition, but only three individuals per condition/sex/stage were analyzed for gene expression. A detailed scheme of the organization of animals per rooms can be found in Additional Fig. 2 and also explained in more detail in the material & methods section. Standard farm routines were followed, such as litter rearrangement to ensure all kits had similar weight, but always according to the experimental criteria. No signs of aggression were detected in any of the experimental conditions tested, likely due to continuous cohabitation since birth [33].

**Fig 2.**
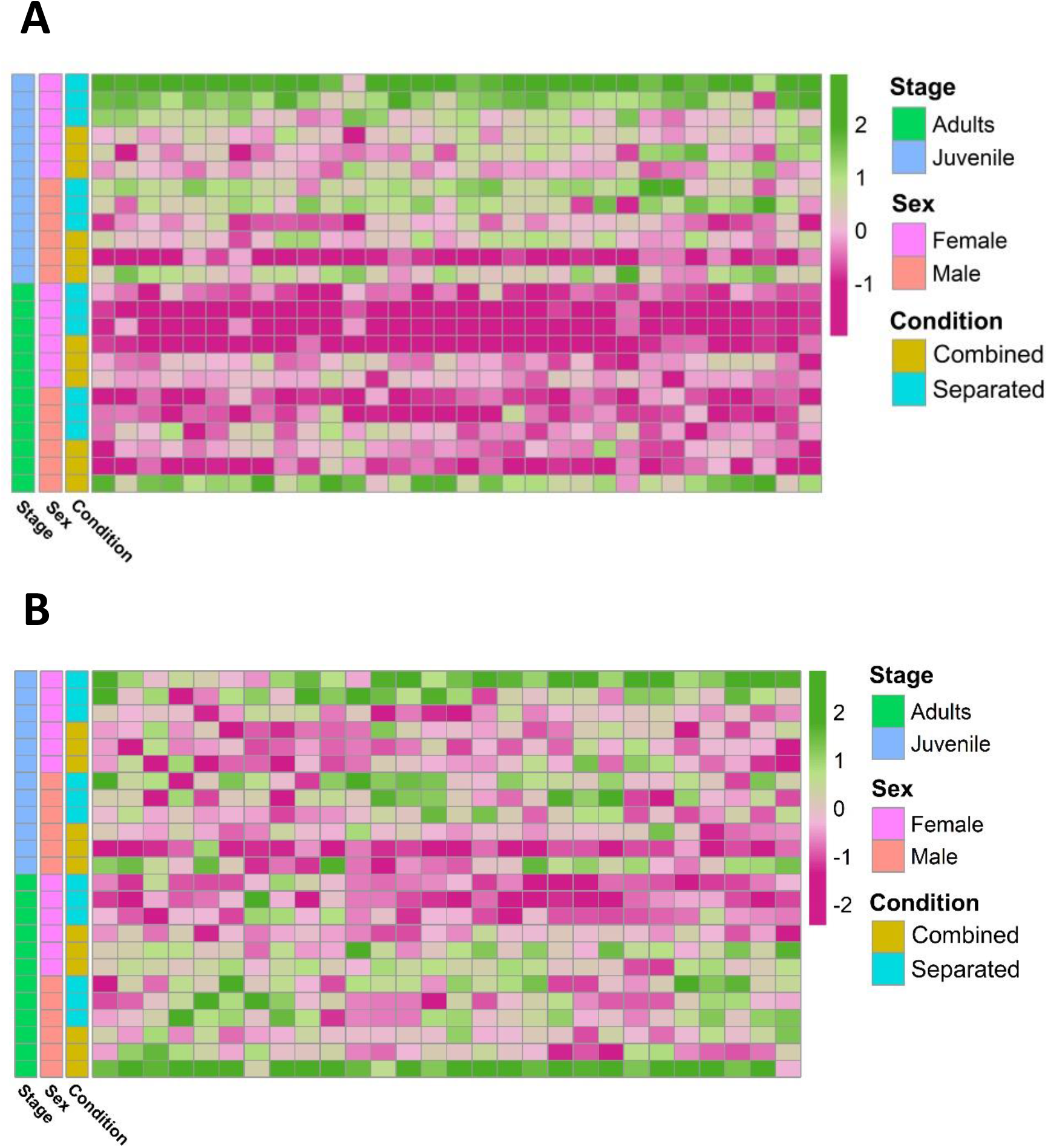
Heatmaps of differentially expressed. **(a)** V2Rs (p value < 0.05) and **(b)** V1Rs (p < 0.05).

Separated female (SF), separated male (SM), combined female (CF) and combined male (CM) groups were constituted both for juvenile (40 days old) and adult (6 months old) rabbits (Fig. 1a; Additional Fig. 1). The two sampling points were selected following Van der Linden et al. [27]; according to these authors, six months would ensure enough time for experience-dependent changes in the abundance of sensory neurons subtypes. Further, we added juvenile individuals to ascertain whether these changes are already occurring at earlier stages. We characterized gene expression in the VNO for the four conditions (SF, SM, CF, CM) and two stages (adults and juveniles) using RNAseq.

### 1. Sex-separation induces sex-specific differences in gene expression of male and female rabbits

We first identified differentially expressed genes (DEG; false discovery rate (FDR) < 0.05) between females and males under sex-separated (SF *vs* SM) or sex-combined conditions (CF *vs* CM) in the rabbit VNO of adults and juveniles (Additional file 1). We found 177 and 66 DEGs, when comparing adults in sex-separated and sex-combined conditions, respectively. The same comparison in juveniles rendered significantly fewer DEGs, but following a similar trend as in adults: 39 DEGs when sexes were separated, while only 2 when sexes were combined (Fig. 1b). These data show that sex-separation determines greater transcriptomic changes between males and females in the rabbit VNO than in the sex-combined scenario, not only in adults but also in juveniles (Fig. 1c,d; Additional Fig. 3). Very few DEGs were shared between both scenarios (sex-separated and sex-combined) in adults (10 genes; Fig. 1b; Additional file 2), demonstrating that the vast majority of DEGs were condition-specific. Seven of those 10 genes were up-regulated in males in both scenarios and included genes such as cholecystokinin (CCK), major allergen I polypeptide chain 1 (FEL1A) or excitatory amino acid transporter 3 (SLC1A1). The latter is a glutamate transporter gene which was previously involved in fear response behaviour, and also obsessive–compulsive disorder [34] and fertility [35] in males.

**Fig 3.**
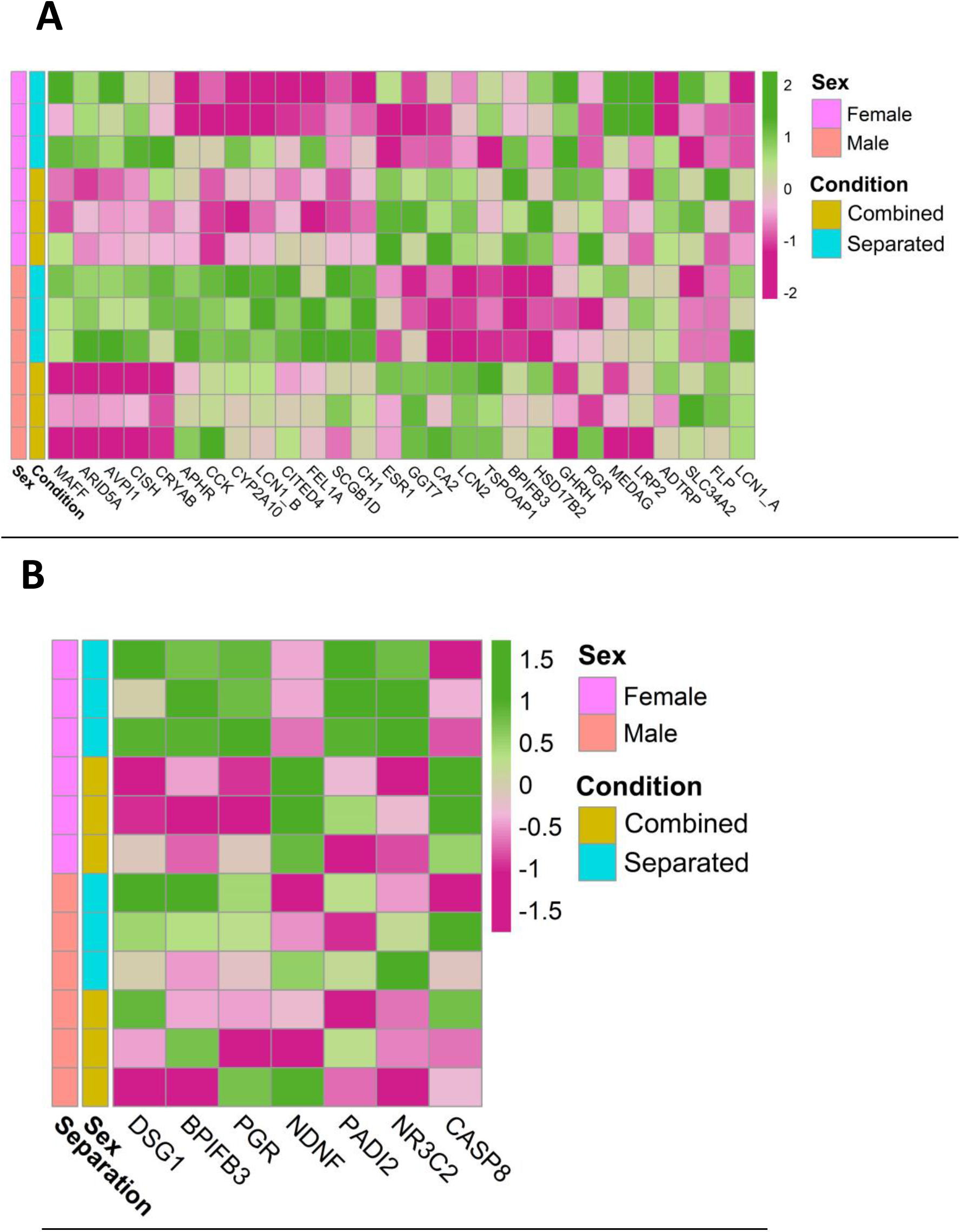
**Heatmap of relevant reproduction-related genes** in **(a)** adult male and female and **(b)** juvenile female rabbit VNO, comparing sex-separated *vs* sex-combined conditions (SF *vs* CF).

In juveniles, all the 39 DEGs found were specific of the sex-separated scenario, suggesting that sex-separation induces much greater differences between sexes at this stage than sex-combined. Only 3 of these DEGs were also detected in the same comparison in adults (Fig. 1b; Additional file 2), therefore, there is almost no overlap between stages. These results suggest that juveniles and adults have sex-specific gene expression repertoires and that both stages should be analyzed independently.

All in all, results indicate that sex-separation induces VNO gene expression differences between males and females in the adult and juvenile rabbit VNO. Adult rabbits exhibit the largest divergence in the number of genes expressed (111 genes) and therefore, a higher plasticity under the two environmental scenarios.

### 2. VNO transcriptome changes depending on environmental conditions in a sex-specific manner

The differential gene expression detected between females and males subjected to different environmental conditions (sex-separated *vs* sex-combined) could arise via changes in one sex or in both sexes. To decipher the influence of each sex, we compared DEGs between sex-separated and sex-combined females (SF *vs* CF) and males (SM *vs* CM), both in adults and juveniles (DEGs combined *vs* separated). In adults, 235 (SF *vs* CF) and 380 (SM *vs* CM) DEGs were detected, while in juveniles the difference was much higher in females (327; SF *vs* CF) than in males (10; SM *vs* CM) (Fig. 1b; Additional Fig. 4; Additional file 3). It may be speculated that the big transcriptomic changes in the VNO of female juveniles could play a role on the onset of female puberty. Moreover, DEGs were mostly sex-specific for each condition, and in adults only 15 DEGs were shared between male and female comparisons (Fig. 1b; Additional file 4). Remarkably, among these 15 DEGs we found the G-protein couple receptor Gαi2, also called GPA1, which is known to induce signal transduction of vomeronasal type-1 receptors (V1R). The fold change (FC) of this gene was among the highest in both sex comparisons (FC: −2.92 in females and −1.91 in males) being up-regulated in the sex-separated condition for both sexes. In juveniles, only two DEGs overlapped between sexes out of 10 DEGs detected in males (Fig. 1b; Additional Fig. 4).

**Fig 4.**
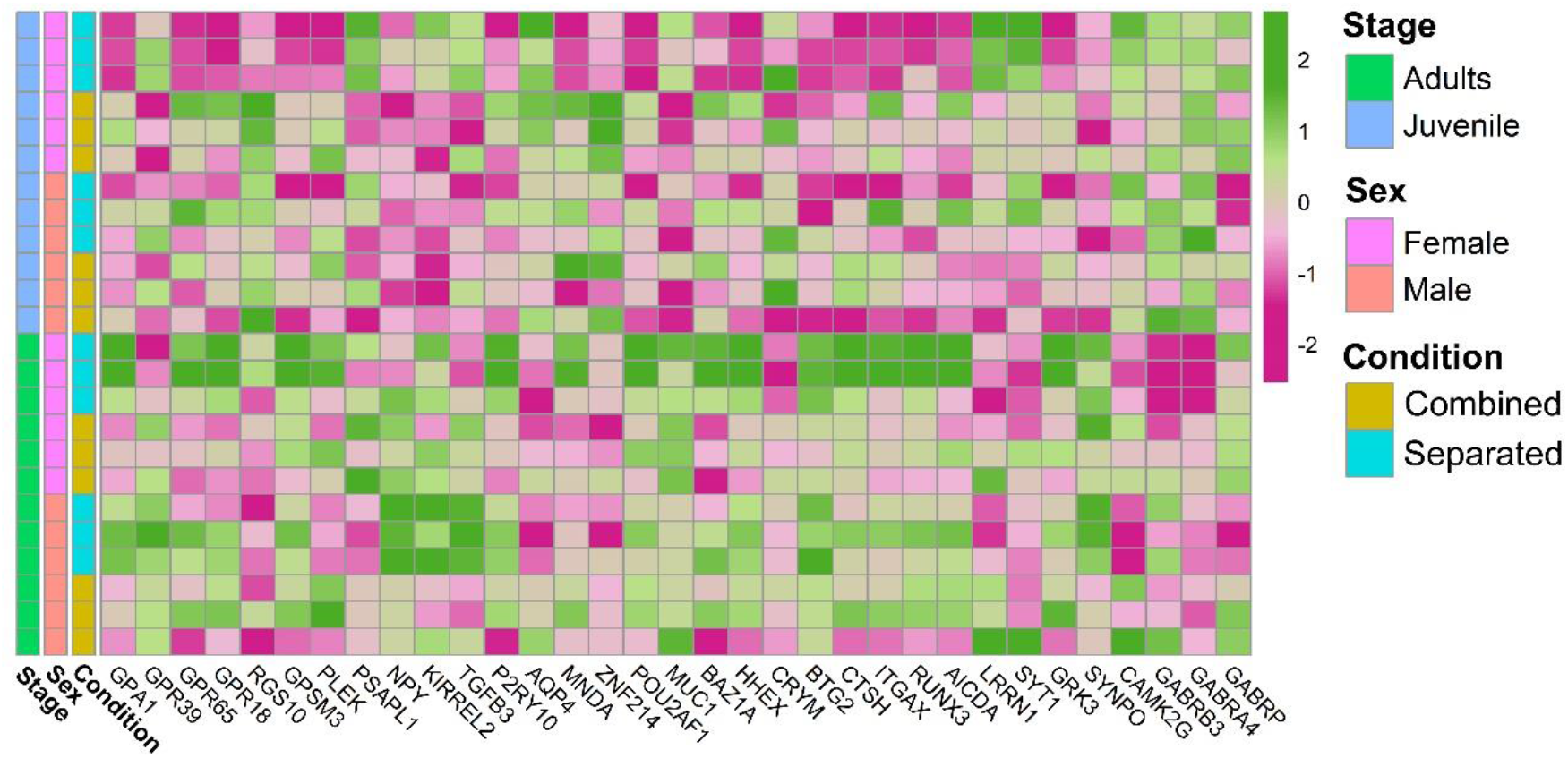
Heatmap of genes related to the VNO functional activity.

A higher overlap was observed when comparing the results of juveniles and adults of the same sex, especially in females, which shared 34 DEGs (Fig 1b; Additional file 4). Gαi2 was shared also in this case, highlighting the importance of this protein not only in adults but also at prepuberal stages; however, unlike the previous comparisons, Gαi2 was up-regulated in the sex-combined condition in females. In males, two genes overlapped between juveniles and adults among the 10 DEGs found in the juvenile comparison (Fig 1b; Additional file 4).

These results indicate that adults of both sexes exhibit specific responses of their overall gene repertoires depending on the environmental conditions (sex-separated and sex-combined scenarios), which is also true for juvenile females. Therefore, when studying how environmental modulation affects VNO gene expression, variations in sex and stage are highly expected and should be carefully considered.

### 3. Vomeronasal receptor repertoire undergoes significant down-regulation in adult female VNO under the sex-separation condition

We specifically analyzed the gene expression of the vomeronasal receptor (VRs) repertoire considering its relevance for chemocommunication, especially on reproductive behaviour. We used a more conservative significance threshold of p < 0.01 (unadjusted) considering the low transcript abundance of VRs, as previously suggested [27]. We also considered as ‘suggestive’ those genes with p < 0.05 (unadjusted) to broadly deepen into the VRs gene repertoire. First, we compared the number of VR DEGs between male and female under sex-separated or sex-combined conditions in adults and juveniles. In adults, we found 4 consistent (p < 0.01; 3 V1R and V2R) and 17 suggestive (p < 0.05; 12 V1R and 5 V2R) differentially expressed (DE) VRs in the sex-separated condition, all of them down-regulated in females (Table 1; Additional file 5). Conversely, only one suggestive DE VR (V1R) was found in the sex-combined condition. These results support the suppression of VRs expression in sex-separated females, while in the sex-combined scenario both sexes showed a similar VR gene expression profile. In juveniles, fewer DE VRs were detected, but their expression followed a similar pattern as in adults: 1 consistent (V1R) and 10 suggestive (3 V1R and 7 V2R) between females and males in the separated condition and none in the sex-combined condition (Table 1; Additional file 5). Contrary to adults, in juveniles we found up-regulation of the ten VRs in females in the sex-separated condition (SF > SM; p < 0.05).

**Table 1.** Differentially expressed VRs among the specific experimental conditions in the VNO of adult and juvenile rabbits and its comparison with previous data in mice VNO. [27] (p < 0.01, consistent difference; in brackets p < 0.05, suggestive difference; (both unadjusted)) SF: separated females; SM: separated males; CF: combined females; CM: combined males.

| Comparisons | Number of DEGs (ADULT) | Number of DEGs (JUVENILE) | Number of DEGs (ADULT MICE) Van der Linden et al. 2018 [27] |
| --- | --- | --- | --- |
| SF vs SM | 4 (17) | 1 (10) | 12 |
| CF vs CM | 0 (1) | 0 | 10 |
| SF vs CF | 3 (12) | 9 (27) | 0 |
| SM vs CM | 2 (8) | 0 (10) | 4 |

We then aimed to evaluate whether the DE VR repertoire is shared between adults and juveniles. We found no overlapping VRs in the four comparisons (SF *vs* SM; CF *vs* CM in both juveniles and adults; p < 0.05), underscoring that the DE VRs are highly sex- and stage-specific, and that the environment critically influences VRs expression. Remarkably, these results contrast with those found by Van der Linden et al. [27] in the mice VNO, where five DE VRs that overlapped between male and female sex-separated (SF *vs* SM) and sex-combined (CF *vs* CM) mice were detected. Accordingly, they concluded that the environment has little effect on VRs gene expression and suggested that differences might be due to self-derived odors. Our results support that each species could show its own VR expression pattern, which undergoes high fluctuation across sexes, stages and species depending on socio-environmental conditions, thus reflecting how dynamic and plastic the VNO must be for external stimuli perception.

We then characterized whether DE VRs detected between female and male rabbit subjected to environment conditions (sex-separated *vs* sex-combined) could originate from changes in one sex or in both sexes. In female adults, we found 3 consistent (V1R, V2R) and 12 suggestive (8 V1R and 4 V2R) DE VRs (SF *vs* CF), whereas in male adults (SM *vs* CM), we detected 2 consistent (1 V1R and 1 V2R) and 8 suggestive (3 V1R and 5 V2R) DE VRs (Table 1; Additional file 6). So, adults of both sexes undergo changes in the expression of VR genes depending on the environmental conditions. Remarkably, most VRs were up-regulated in the sex-combined condition (CF > SF; CM > SM), which indicates that sex-separation does not promote VRs expression in the adult rabbit VNO, especially in females. Additionally, there was no overlap between sex comparisons (SF *vs* CF and SM *vs* CM), suggesting sex-specific gene expression repertoires under environmental modulation. In juveniles, we found 9 consistent (1 V1R and 8 V2R) and 27 suggestive (6 V1R and 21 V2R) DE VRs between sex-separated and sex-combined females (SF *vs* CF), whereas we detected no consistent and 10 suggestive (5 V1R and 5 V2R) DE VRs for the same comparison in males (SM *vs* CM) (Table 1; Additional file 6). Contrary to adults, in juveniles all VRs were up-regulated in sex-separated animals both in males and females. We found four genes overlapping between males and females for those comparisons (SF *vs* CF and SM *vs* CM), suggesting similar environmental influence on those genes for both sexes.

Two main conclusions can be extracted from the VRs gene expression results: 1) VRs gene expression is sex- and stage- specific under different socio-environmental conditions (separated *vs* combined) and 2) adult females in a sex-separated scenario showed the lowest VRs expression among all comparisons. For a more refined evaluation of VRs gene expression, we then visualized all V2R and V1R showing differential expression (consistent or suggestive) in all comparisons as a heatmap (Figs. 2a,b). V2Rs expression was significantly higher in juveniles than in adults regardless of sex and environment, and the lowest V2R expression was shown by sex-separated adult females. Interestingly, one sex-combined adult male and one sex-separated juvenile female showed specific up-regulation of V2Rs compared to the rest of individuals (outliers), hampering significant conclusions due to the increased variation (Fig. 2a). The pattern of V1R expression underwent greater interindividual variation than V2R, but still showed that sex-separated adult females display the lowest V1Rs expression (Fig. 2b) - although not so extreme as for the V2R expression. Furthermore, the same two outlier individuals for V2Rs also showed higher overall V1Rs expression. The high interindividual variation raises the question of whether these two animals could be dominant individuals, being more prepared than others for mating or fighting purposes.

Individual variation of VRs expression has been previously described [36], but most VRs studies have emphasized results per group of individuals under the same experimental conditions [27]. Our data show significant intragroup variation in the VRs gene expression repertoire despite the similar conditions assessed (sex, stage and environment), highlighting the importance of individual variability that makes difficult extracting consistent conclusions between groups. Further VRs studies should always consider this interindividual variability trying to minimize error in the estimations, which means randomizing genetic variation, homogenizing as much as possible environmental conditions and increasing sample size.

### 4. Environmental modulation triggers differential expression of VNO genes involved in reproduction and sexual behaviour, supporting its plastic capacity to ensure individual survival

Adult males and females exhibited sex-specific expression differences of reproduction-related genes, subjected to environmental conditions. Genes belonging to the lipocalin family followed significant differential expression according to conditions in males and females (Fig. 3a) (e. g. Lipocalin 1A (LCN1_A, ENSOCUG00000026763), SM > SF; lipocalin 1B (LCN1_B, ENSOCUG00000028189), SM > CM; lipocalin 2 (LCN2, ENSOCUG00000021002), CF > SF, CM > SM, SF > SM; and aphrodisin (APHR, ENSOCUG00000029236), SM > SF)), suggesting the specificity of different members of the lipocalin family, which may bind different putative sex-specific pheromones depending on the environmental conditions, which in turn would have an impact on the VNO pheromone-detection. We also found differential expression of BPIFB3 (ENSOCUG00000014883; CM > SM), a gene belonging to the BPI family that interacts with a major urinary protein -MUP10- in mice, a lipocalin binding pheromone protein affecting the sexual behaviour of females (https://string-db.org/network/10090.ENSMUSP00000095655). A similar expression pattern (CM > SM) was found for the female-specific lacrimal gland protein (FLP, ENSOCUG00000027780), a gene previously identified as a lipocalin in hamsters which shows 85% protein sequence identity with male-specific submandibular salivary gland proteins (MSP) secreted in saliva and urine of male hamsters [37]. Therefore, rabbit FLP may be other lipocalin subjected to environmental modulation, whose presence in the VNO points to a role as carrier of small putative hydrophobic pheromones. Further studies are required to determine the role of each lipocalin within the rabbit VNO. We also found differential expression of other important reproduction-related genes in adults, such as the sex-steroid binding progesterone receptor (PRG, ENSOCUG00000014693) and estrogen receptor (ESR, ENSOCUG00000004829), both CF > SF, p = 0.057 (adjusted) and p = 0.014 (unadjusted), respectively. Other genes related to steroid-binding (FEL1A, CH1 CITED4, CISH, etc.) showed differential gene expression between conditions (Fig. 3a; Additional Table 1). In juveniles, most of the analyzed reproductive-related genes were found down-regulated compared to adults (Additional Fig. 5; Additional Table 1), probably because they have not reached puberty yet. Additionally, females but not males showed significant differences between separated and combined conditions (SF *vs* CF). Reproductive-related genes, such as BPIFB3, DSG1 (desmoglein 1, which responds to progesterone) and PGR (this latter, p < 0.05 (unadjusted)), were differentially expressed between conditions (SF > CF) (Fig. 3b; Additional file 3), suggesting that gene expression changes of reproduction-related genes occur at least from day 40 in the female VNO. Interestingly, NDNF (neuron derived neurotrophic factor, ENSOCUG00000002038), a gene directly related to GnRH neuron migration and the onset of puberty [38,39,40], was significantly up-regulated in combined females compared to separated females (CF > SF) (Fig. 3b; Additional file 3), and therefore exposure to different environments may influence the timing of puberty. We previously showed that rabbit juveniles already expressed genes related to reproductive functions [32], and here we have proved that their expression is modulated under different environmental conditions.

**Fig 5.**
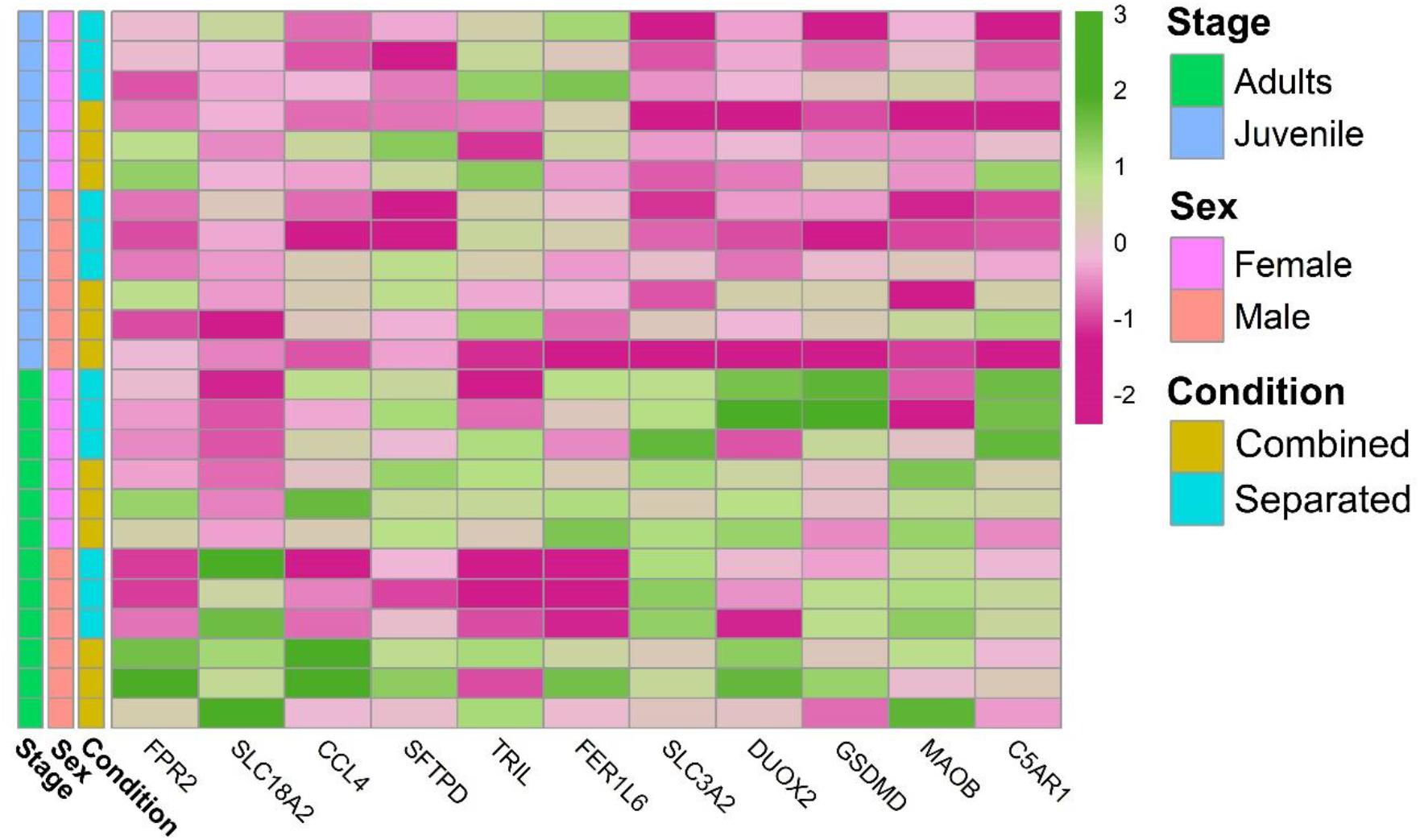
Heatmap of genes related to the VNO capability of detecting external stimuli, harmful pathogens and its relation with the immune system.

Taken together, these results indicate that, in the adult and juvenile VNO, expression of several reproduction-related genes changes according to the environmental conditions (sex-separated, sex-combined) and some of them are sex-specific.

### 5. Environmental changes modulate VNO functional activity, thus adding complexity and flexibility to the VNO sensory code

We next sought to determine whether the functional activity of the VNO varies among the experimental groups. Enrichment analyses revealed overrepresentation of G-protein related terms such as ‘G protein-coupled chemo attractant receptor activity’ and ‘G protein-coupled receptor signaling pathway’ among DEGs detected in separated *vs* combined females (SF *vs* CF), both in adults and juveniles (Additional file 7). In juveniles, these terms as well as others related to ‘response to stimulus’ and ‘signal transduction’ were directly linked to vomeronasal type-2 receptors and the G protein GPA1 (alpha subunit of the heterotrimeric guanine nucleotide-binding G protein), which is known to mediate mating pheromone signal transduction in yeast and is found expressed in V1R and V2R expressing neurons in mammals [41]. Both in juveniles and adults GPA1 was found up-regulated in separated females when compared to combined females (SF > CF), and a similar pattern was found in the adult male comparison (SM > CM). Interestingly, a heatmap representation (Fig. 4) shows that GPA1 is more expressed in adult separated females than in any other group. It is known that sensory transduction through the VNO proceeds via a G-protein-coupled mechanism [42]. Our data showed that a high number of G-protein coupled receptor genes (GPR39, GPR65, GPR18), as well as genes which modulate (RGS10, GPSM3, PLEK) or inhibit (PSAPL1) the G-protein signaling activity, are differentially expressed, especially when comparing sex-separated and sex-combined female adults (SF *vs* CF) (Additional File 3; Fig. 4). We also detected several genes previously involved in VNS neural activity (Neuropeptide Y and Kirrel2) [43, 44] and signal transduction (G proteins such as P2RY10) up-regulated among DEGs in several comparisons (Additional file 3; Fig. 4). These results indicate that G-protein expression is especially up-regulated in separated adult females, but its expression varies among comparisons, likely cooperating in the modulation of the vomeronasal signal transduction in response to specific environmental inputs.

Another gene family that might be related to VNO signal transduction and worth mentioning is water channel aquaporins (AQPs). In the rabbit VNO transcriptome, we previously detected high expression of AQP2, AQP3, AQP4 [32]. Other studies proved the expression of AQP2 and AQP3 in mice olfactory organs but not in their sensory neurons [45], whereas AQP4 was the only aquaporine substantially expressed in VNO sensory neurons in rat [46]. Remarkably, we detected AQP4 down-regulated in the VNO of adult male rabbits when sex-separated compared to sex-combined (CM > SM) (Additional file 3; Fig. 4), thus suggesting that AQP4 undergoes gene expression changes in a sex-specific manner, yet their presence in VSNs in rabbits remains unknown. AQP4 might be involved in social behavior and sexual preference by having a specific role in the VNO neuronal transduction. However, its physiological importance in sensory neurons of the VNO remains to be discovered [45].

Other enriched functions were detected under environmental changes (Additional file 7). We found down-regulation of biological processes GO terms such as ‘gene expression’ and ‘regulation of transcription’ in the list of DEGs between adult separated females *vs* separated males (SF *vs* SM). In this comparison as well as in others such as separated females *vs* combined females (SF *vs* CF), both in juveniles and adults, we also found down-regulation of terms related to ‘RNA metabolic processes’. Among the genes involved in those functions, we found several transcription factors (MNDA, ZNF214, POU2AF1, MUC1), transcript repressors (BAZ1A, HHEX, CRYM, BTG2), modulators of gene expression (CTSH, ITGAX, positive regulation; and SERPINF1, negative regulation) and others related to epigenetic changes (RUNX3, AICDA). More specifically, genes involved in the regulation of synaptic transmission (LRRN1, SYT1, GRK3) as well as in synaptic plasticity (SYNPO, CAMK2G) and GABAergic activity (GABRB3 and GABRA4, GABRP) were also DE among comparisons (Additional file 3; Fig. 4), thus supporting the VNO functional plasticity. All these genes might play a role in how sensory inputs are perceived by VNO –likely by modulating the expression of vomeronasal receptors– and how the electrical responses are sent to higher brain centers.

These results support that the VNO undergoes transcriptional changes according to environmental conditions, especially down-regulating regulatory and transmission elements in separated adult females, which might represent an adaptation of sensory responses to upcoming signals. These results are consistent with recent data by Flores-Horgue et al. [30] in the main olfactory epithelium, in which odorant exposure led to reprogramming via the modulation of transcription.

### 6. The VNO as a first barrier in the detection of external stimuli, including harmful pathogens, and its relation with the immune system

The VNO is capable of detecting a broad range of chemical cues which fluctuate depending on given environmental scenarios. We found enriched molecular function GO terms related to ‘response to chemical/biotic stimuli and to other organisms’ and ‘regulation of response to external stimulus’, especially in separated *vs* combined females (SF *vs* CF), both in juveniles and adults, but also in separated female *vs* separated male (SF *vs* SM) in adults. The VNO is provided with different types of vomeronasal receptors to detect those stimuli. In addition to V1R and V2R receptors, several H2-Mv –related to ultrasensitive chemodetection– and FPRs – linked to pathogen sensing– genes have been identified in mice [10,12,13]. We blasted the nine mice H2-Mv receptors against the rabbit VNO transcriptome [32] and found significant hits with six genes belonging to the class I MHC. Among them, two showed gene expression variation depending on sex-combined *vs* sex-separated scenarios (Additional file 8). These results suggest that different vomeronasal receptors belonging to the MHC class I receptors previously unidentified in rabbits might play a role as vomeronasal receptors with plastic capacity to sensitively detect specific external cues in the rabbit VNO.

The VNO has also been involved in the detection of harmful bacteria and sick conspecifics, likely through FPRs [18]. The rabbit genome does not contain expansion of VNO FPRs, but the rabbit VNO transcriptome includes the immune receptors FPR2 and FPR1 [32]. Here we found up-regulation of FPR2 and FPR1 in sex-combined individuals, especially in males. Specifically, FPR2 (ENSOCUG00000008606) was up-regulated in adult males when sex-combined, compared to sex-separated males (CM > SM), but also to sex-combined females (CM > CF), this latter without reaching statistical significance (p < 0.05 (unadjusted)). Additionally, FPR1 (ENSOCUG00000008600) was up-regulated in combined males and females when compared to their separated counter partners (CM > SM; CF > SF, for both p < 0.01 (unadjusted)). These results indicate that FPR2 and FPR1 are environment-modulated and therefore its immune function could be more complex than previously thought. Additionally, we found many molecular function GO terms related to ‘response to bacterium’ and ‘defense response’ among DEGs in several comparisons (Additional file 7). Specifically, some DEGs were related to response to toxic substances (SLC18A2, CCL4), defense response to bacteria (SFTPD, TRIL, FER1L6) and virus (SLC3A2, DUOX2), reaction to danger signals (GSDMD and MAOB), or showed specific defense response to Gram-positive bacteria (C5AR1, also known to detect sensory perception of chemical stimulus). Other GO terms directly related to the immune system such as ‘antigen binding’, ‘immune response’, ‘MHC protein complex’ were also found among comparisons (Additional file 7) as well as genes related to the immune system such as immunoglobulins, interleukines and chemokines (Fig. 5; Additional file 3).

Overall, these results indicate that the VNO plays an important plastic role at detecting chemical stimuli and harmful bacteria, in narrow connection with the immune system, thus acting as an interface between the external world and the central nervous system for modulating responses against potential dangers. However, despite the clear importance of the VNO in the detection of harmful stimuli, we are still far from having a clear picture of species-specific VNO receptors in charge of mediating these fundamental behaviours.

## Discussion

We have shown that VNO gene expression in rabbits is exacerbated when there is no contact between individuals of the opposite sex. The low overlap between the different comparisons suggests that variation in the VNO gene expression under environmental conditions is both sex- and stage-specific.

### 1. VNO and VR repertoires show species-specific and environmentally modulated expression: a comparison between rabbits and mice

Our results proved that sex-separation induces sex-specific differences in the rabbit VNO gene expression, and this is consistent with previous results in adult mice -106 and 64 DEGs for the SF *vs* SM and CF *vs* CM, respectively- [27]. However, some features are species-specific. In adult rabbits we detected more DEGs between separated and combined males (SM *vs* CM – 380 DEGs) than in the same comparison of females (SF *vs* CF – 235 DEGs), even though both sexes underwent important transcriptomic changes. Conversely, we detected in juveniles much more DEGs in females (SF *vs* CF – 327 DEGs) than in males (SM *vs* CM – 10 DEGs). Van der Linden et al. [27] reported that females (SF *vs* CF – 64 DEGs) exhibit more changes than males (SM *vs* CM – 35 DEGs) in the adult mouse VNO. Therefore, juvenile rabbits follow a similar pattern to that of adult mice (SF *vs* CF > SM *vs* CM), whereas adult rabbit males showed higher expression differences than females, although these were notable in both sexes. In rabbits, there were hardly DEGs overlapping between sexes (6.38% in adults, 15 of 235 DEGs from SF *vs* CF were also found in SM *vs* CM; and 0.61% in juveniles, 2 of 327 DEGs from SF *vs* CF were also found in SM *vs* CM), in contrast with mice, which exhibited a much higher overlap between SF *vs* CF and SM *vs* CM (28.13%, 18 of 64 DEGs; see Fig. 1g from Van der Linden et al. [27]). These data indicates that the VNO transcriptome is more sex-specific in rabbits than in mice when sex-separated and sex-combined conditions are tested. The fact that mice were laboratory domesticated animals, while rabbits were under farm conditions, might be connected to this observation, e.g. reduced plasticity in laboratory animals.

When comparing gene expression of the VNO adult males and females, we detected a rather small overlap between sex-separation and sex-combined scenarios (15.2% DEGs, 10 of 66 DEGs from CF *vs* CM were also found in SF *vs* SM). In the mice VNO, Van der Linden et al. found much higher overlap (43.8%; 28 of 64 DEGs) between the same comparisons (see Supplementary fig. 6a in [27]. When studying the main olfactory epithelium (MOE), they found again an important overlap (39% (16 of 41 DEGs; see Fig. 2a in [27]) between the same comparisons. However, authors argued little overlap in the adult mice MOE despite both percentages −43.8% and 39%- are rather close, suggesting caution on this conclusion.

When considering specifically the VRs, we could not find overlap between sex-combined and sex-separation scenarios (CF *vs* CM and SF *vs* SM), both in adult and juvenile rabbits. However, Van der Linden et al. [27] found significant overlap between the two comparisons (50%, 5 of 10 VRs in CF *vs* CM were also found in SF *vs* SM; see Supplementary fig. 6e in [27]). They argued that differences in the expression of VRs would be largely independent from socio-environment conditions and may instead be driven primarily by self-derived odors. However, the same comparison for olfactory receptors (ORs) found no overlap between comparisons - similar to what we found in the rabbit VRs-, concluding that the environment would be critical for sex-biased ORs expression. Our data indicates that VRs expression repertoire in rabbits is environmentally modulated and the conclusions obtained by Van den Linden et al. [27] in mice do not apply to rabbits. Therefore, the detection of both self- and opposite sex- derived odors in rabbits could be mediated by either VRs or ORs depending on the perceived stimuli. Our results also highlight that generalized statements across species may lead to mistaken conclusions.

Additionally, Van der Linden et al. [27] studied the expression of 6 VRs using RNA-FISH and only two of them showed consistent profiles with the RNAseq data. They argued that some VR expression differences observed by RNAseq may reflect differences in cellular VR transcript levels and not cell density. However, FISH might not be a suitable approach to characterize VRs plasticity under environment-modulated conditions, especially due to the high sequence similarity shared by different VRs. Instead, new –OMICS techniques such as single-cell and spatial transcriptomics could provide a more solid framework not only on VRs plasticity studies but also on a broader range of vomeronasal receptors and other VNO-related genes, to validate the correlation between transcript abundance and cell density depending on the environmental conditions.

All in all, we found little DEGs overlap in the rabbit VNO between males and females either juveniles or adults in all scenarios tested, which indicates that environmental modulation changes the VNO gene expression in a sex- and stage- specific manner in this species. This leads us to conclude that sex-separation is a condition which induces sex- and stage- VNO specific differences, but also depends on species (mouse *vs* rabbit) tissues (vomeronasal *vs* main olfactory organs) and individuals. Our observations are consistent with the plastic capacity of the VNO to cope with different environmental scenarios, but further studies are needed to clarify the rationale behind such heterogeneity.

### 2. The lipocalin aphrodisin is up-regulated in sex-separated adult male VNO

Different members of the lipocalin family are involved in pheromone transport and can be found in a variety of body fluids such as urine [47], tears [48] or saliva [49]. They can act either as carrier proteins for small hydrophobic like-pheromones -some of them expressed in the VNO [50]- or directly as pheromones, such as darcin in mice [51]. APHR, a pheromone that belongs to the lipocalin family, is known to be released by hamster vaginal secretions (even before the onset of puberty) and parotid gland [52, 53] and it is detected by the male vomeronasal system eliciting copulatory behavior in male hamster [54–57]. Different pheromone-like hydrophobic compounds were specifically bound onto natural APHR suggesting its role as pheromone transporter [57, 58].

We found for the first time expression of APHR in the VNO tissue of rabbits, being highly-expressed in sex-separated males (SM > SF, p < 0.05 (adjusted)). Therefore, its expression appears to be sex-specific and clearly depends on the environment. This could prepare separated males to be ready (receptors not saturated) to bind female-pheromones, eliciting copulatory behaviour in the presence of females. Also, it could selectively modulate the activation of vomeronasal receptors. Interestingly, when both sexes are reared together, the expression of this gene is lower (CM < SM), p = 0.013 (unadjusted)), which could reflect saturation when males are continually exposed to the putative female pheromones. This would follow a similar rationale of the theory described for VRs ‘use it and lose it’ [27], which could then apply to other genes directly involved in pheromone-sensing. Of note, to date lipocalins were only found in the glandular region and vomeronasal mucosa of the VNO, but further studies are needed to decipher where rabbit lipocalins are located within the VNO.

### 3. Sex-steroid receptors are modulated by the environment and may arise as a new type of VNO pheromone receptors

Steroids are well-known for their essential role in animal physiology, particularly regarding the internal state of the individual. Less studied but also vital for animal behaviour is the role of steroids in chemosensation. These compounds are excreted by animals in urine or faeces, releasing a readout of its own biological state, and therefore acting as chemosignals/pheromones to be detected through the VNO of conspecifics [21]. Specifically, estrogens and oxytocin have proved to modulate VNO neural excitability inducing behavioural changes [59, 60]. VSNs are highly sensitive to excreted glucocorticoids, sex-steroids and bile acids that might act as pheromones [21]. Indeed, sulfated compounds are the predominant VSNs ligands in female mice urine [61]. Therefore, sex-hormones play an important role in activating VSNs, modulating sensory signaling and contributing to the heterogeneity of VSNs response capabilities [62, 63]. However, the relationship between steroid cues and the receptors involved in its detection, as well as the neural pathway involved in behavioural response remain still unclear. Recently, it has been shown that V1R receptors are involved in the detection of sulfated estrogens [8] and bile acids [64]. These are likely the predominant receptors of steroids in mice, but V2Rs also seem to be involved [21]. Noteworthily, sex-steroid receptors were found in VSNs, suggesting a potential role in signaling actions of pheromones [65]. For instance, estrogen receptor (ESR1) in the VNO is directly localized to neural progenitors and mediates neurogenesis during pregnancy in female mice [66]. In the rabbit VNO, we found progesterone and estrogen receptors as well as other sex-steroid binding molecules, whose expression pattern is modulated by specific environmental conditions. This highlights the plastic capacity of sex-steroid receptors in the rabbit VNO, placing them as potential candidates to perceive particular external and internal cues and trigger specific signaling responses.

The complexity of steroid chemosensation suggests that there is not a universal rule governing sex-steroid pheromone *vs* receptor activity [21]. However, throughout evolution, receptors of different families may have acquired steroid sensitivity. We argue that sex-steroid and VR receptors in the VNO may be responsible for detecting sex-steroid cues and triggering signal transduction to brain centers. We propose two possible hypotheses: 1) sex-steroid receptors may act as a first screening of sensory perception in response to specific internal states followed by the activation of VRs of VSNs. Accordingly, we would expect VRs and sex-steroid receptors to be found in the same VSNs, to avoid energy loss due to cell-to-cell communication as well as to maintain high efficiency to trigger a quick response; and 2) both types of receptors, VRs and sex-steroid receptors perceive pheromones independently. In this case, we would expect VRs and sex-steroid receptors to be found in different VSNs and, accordingly, sex-steroid receptors could be considered as a new type of pheromone-receptors. Consequently, understanding the cellular rationale of the VNO will provide the needed framework for deciphering how pheromonal inputs are mediated by sensory neurons, potentially placing VSNs as a peripheral structure that integrates internal states with external chemosignals.

When studying chemosensation, another layer of complexity needs to be added. On the one hand, steroid-binding globulins have been found in VNO mucus but not in VSNs, and could sequester additional pheromones to deliver them to sensory cells –similar to the role of lipocalins– [67]. Additionally, VNO is heavily vascularized, especially in rabbits [68], and therefore accessible to self-circulating steroid hormones and peptides, and thus an individual could react to their own steroids. Indeed, mice can regulate sensory signal depending on their own hormonal state [69]. Furthermore, estradiol can be locally synthetized in the VNO modulating odorant responses mediated by VSNs, probably through estrogen receptors found in VSNs [70]. Finally, in some cases, the integrated action of various chemical compounds is needed (i.e. urinary estrus signals and sulfated estrogen) to trigger a specific behaviour [71].

### 4. A potential close relationship between the VNO and female puberty

Females exposed to urine odors of males during prepubertal period accelerate puberty, a phenomenon known as Vandenbergh effect [72]. Removal of either both olfactory bulbs or one olfactory bulb and sectioning the vomeronasal nerve abolished the rapid increase in uterine weight of immature female mice exposed to male mice, likely through pheromone-mediated behaviour [73]. Separated females that are exposed to male odours around the puberty period showed an increase in c-Fos activity (neuron marker activity) in the accessory olfactory bulb and higher centers of the vomeronasal pathway, arguing that such activity might be related to an earlier puberty onset [74]. Additionally, it has been suggested that lesions of the VNO will block puberty acceleration in female mice exposed to male mouse urine [74]. A recent study has determined that peripubertal VNO ablation decreased sexual odor preferences and neural activity in response to opposite-sex odors, and drastically reduced territorial aggression in male mice, suggesting that the VNO contributes to the sexual differentiation of behavior and neural response to conspecific odors [75].

Our results indicate that the VNO of female juveniles shows 327 DEGs between sex-separated and sex-combined conditions (SF *vs* CF), whereas only 10 DEGs were found in the same comparison in males (SM *vs* CM). These differences highlight the plasticity of the VNO in juvenile females exposed to different conditions, which might be linked to female puberty. Regarding VRs, males and females showed similar VR repertoire when sex-combined (CM *vs* CF), whereas 10 DE VRs were found differentially expressed when sex-separated (SM *vs* SF), all of them up-regulated in females. Importantly, all 27 DE VRs found between separated and combined females (SF *vs* CF) were up-regulated in separated individuals. We speculate that this VR up-regulation in juvenile separated females might follow the theory ‘use it and lose it’ [24, 27], suggesting that females not exposed to males would show standard VRs expression in a given VSN; however, after even short male odor stimulation, which is known to accelerate puberty, VRs expression would increase until a certain threshold, followed by selectively reduction of VRs expression and possibly in the abundance of specific VSN subtypes as well. In other words, pheromones contained in male urine would bind already activated VRs of separated females, and this would accelerate puberty onset; therefore, puberty onset would depend on the environmental condition and the sex-separated scenario would ‘prepare’ VRs to receive pheromones from the opposite sex. Our data suggest that this interpretation would be only applicable for juveniles and puberty onset. VR gene expression proved to be down-regulated in separated adult females, which would therefore follow a complete different rule. One suggestion is the theory ‘use it or lose it’ [24, 27]. This means that females that are not exposed to males for a long period of time (6 months) might reduce their capacity to respond to male cues. Of note, down-regulation of VRs in separated females contrasts with the up-regulation of GPA1 in separated females and also in separated males, a G-protein known to mediate V1R and V2R signal transduction, raising the question of whether GPA1 could be implicated in other G protein coupled receptor pathways and therefore modulate VNO signaling in response to other vomeronasal receptors.

## Conclusions

This study shows the plastic capacity of the rabbit VNO, and that in mammals this feature is not limited to mice. Our experimental design of sex-separation *vs* sex-combined scenarios is a suitable model of environmental modulation to study VNO plasticity. We demonstrated that the rabbit VNO exhibits significant gene expression differences between male and female under sex-separation but not sex-combined scenarios. These differences are not only remarkable for the vomeronasal receptor repertoire but also for many genes involved in sensory perception, reproduction, immunity and VNO functional activity.

Regulation of transcription arises as a mechanism that the VNO may use to adapt sensory responses to specific external stimuli. Further studies should approach the molecular logic of such transcriptional changes as well as its possible relationship with variation in the VNO cellular and neuron abundance.

## MATERIAL & METHODS

### Experimental design

All animals were reared under the same farm environment and management protocols. For the experiment, we employed 1) male and female rabbits that were separated at birth (sex-separated) so that males and females did not have contact with members of the opposite sex, and compared them to 2) male and female rabbits that were in contact (sex-combined). Two experimental time points –juveniles (40 days) and adults (6 months)– were considered. Separated female (SF), separated male (SM), combined female (CF) and combined male (CM) groups were constituted both for juvenile (40 days old) and adult (6 months old) sampling (Fig. 1a; Additional Fig. 1).

Four different rooms were employed in the experiment: room a, c, d for sex-separated conditions, and room b for sex-combined condition. All rooms had three double rows with capacity for 280 mothers with their litters per row (a total of 840 mothers/room) (Additional Fig. 2). For the sex-separation condition, we first employed room a, in which male and female litters were placed in different rows in opposite sides of the room to avoid any type of contact between sexes. Weaning took place on day 30, mothers were then taken out of the boxes while kits remained until day 40 (sampling point 1: juveniles). 60 males and 60 females of 40 days old were then separated into different rooms (room c and d for males and females, respectively) of the farm. We kept four animals/cage and used 15 cages (60 rabbit males in room c and 60 rabbit females in room d), to ensure the same environment until six months old (sampling point 2: adults). For the sex-combined group we used a different room, room b, in which four kits of each sex were cared per mum (8 in total). We arranged 100 mothers, each with their own litter to ensure exposure to the same environment. The protocol was similar as for the sex-separated scenario, but no changes between different rooms were performed. Of note, since males and females were kept together to ensure direct contact between individuals, kits born during the six months of the experiment were removed from the box on day 0 to avoid exposure to pup-specific odors (same criteria as Van der Linden et al. [27]).

### Sampling

24 animals, separated in 12 juveniles and 12 adults of 40 and 180 days, respectively, were used including the same number of males and females within each group. All animals pertained to a commercial hybrid -Hyplus strains PS19 and PS40 for female and male, respectively- and were maintained on a farm (COGAL SL, Rodeiro, Spain) under the same temperature conditions (18-24°C), dark-light cycles of 12:12 hours and *ad libitum* feeding and drinking. All individuals were humanely sacrificed by an abattoir of the same company, following strict ethical farm conditions and in accordance with the current legislation. Sex-separated animals were sacrificed at the beginning of the day to avoid any type of contact with members of the opposite sex, and males and females were sacrificed in consecutive different days for the same reason. Their heads were separated from the carcasses in the slaughtering line and the double VNO structure was immediately dissected out after opening the lateral walls of the nasal cavity and removing the palate and nasal turbinates. Samples were immersed in Trizol and kept in ice (∼4°C). Tissue was homogenized at the sampling point using a mixer to guarantee the whole tissue sample was soaked by Trizol-due to the double bone and cartilage envelope of the rabbit VNO-. After 20 minutes, all samples were stored at −80°C for further RNA extraction.

### Transcriptomic Analysis

RNA extraction, library preparation and bioinformatic analysis was previously described in [32]. Briefly, libraries were sequenced on an Illumina Nova-Seq 6000 150 bp PE run by Novogene. The quality assessment and the filtering and removal of residual adaptor sequences were performed using FastQC v.0.11.7 (https://www.bioinformatics.babraham.ac.uk/projects/fastqc/) and on Trimmomatic v.0.39 [76], respectively. Filtered reads were aligned against the rabbit genome (OryCun2.0; [77] and assigned to genes based on the latest annotation of the rabbit genome [78] using STAR v.2.7.0e [79] two-pass mode.

Gene count data were used to calculate gene expression and estimate differential expression (DE) using the Bioconductor package DESeq2 v.1.28.1 [80] in R v.3.6.2 [81]. Size factors were calculated for each sample using the ‘median of ratios’ method and count data were normalized to account for differences in library depth. Normalized reads were used as a measure of expression and were calculated by taking the average of the normalized count values, dividing by size factors, taken over all samples. This corresponds to the ‘basemean’ obtained with DESeq2. Next, gene-wise dispersion estimates were fitted to the mean intensity using a parametric model and reduced towards the expected dispersion values. Finally, differential gene expression was evaluated using a negative binomial model that was fitted for each gene, and the significance of the coefficients was assessed using the Wald test. The Benjamini-Hochberg FDR correction for multiple tests was applied, and transcripts with FDR < 0.05 were considered DEGs. Hierarchical clustering and PCA were performed to assess the clustering of the samples and identify potential outliers over the general gene expression background. The R packages “pheatmap”, “PCAtools”, “EnhancedVolcano” and “UpSetR” were used to plot heatmaps, principal component analysis (PCA), volcano plots, and the upsetplot, respectively. Gene functional annotation was performed by searching all rabbit VNO GO terms with PANNZER2 [82] and then assessing the enriched terms for each comparison of our study (SF *vs* SM, CF *vs* CM, SF *vs* CF, SM *vs* CM) and also for each condition independently (SF, SM, CF and CM) both in juveniles and adults, by using AgriGO v2.0 [83]. Gene BLAST analyses were performed using Ensembl BLASTN tool (https://www.ensembl.org/Multi/Tools/Blast).

In parallel, we also performed our analysis using kallisto [84], which pseudoaligns reads to a reference, producing a list of transcripts that are compatible with each read while avoiding alignment of individual bases, which is what STAR does. Similar results were obtained with both software, STAR and kallisto, and therefore STAR was used by default in our analysis.

## Additional data

**Additional Figure 1.**
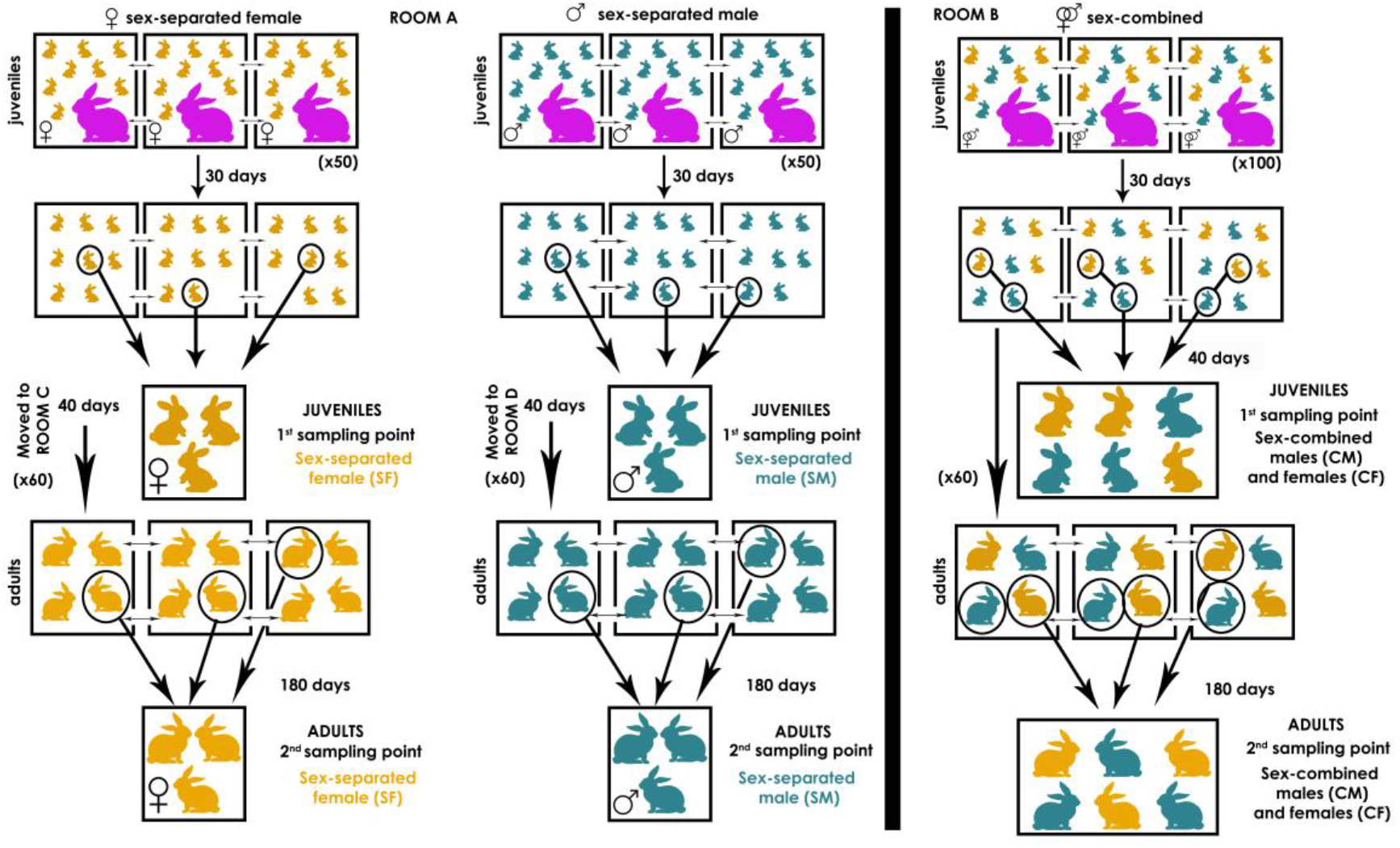
Experimental design of sex-separation and sex-combined male and female, both for juvenile and adult individuals

**Additional Figure 2.**
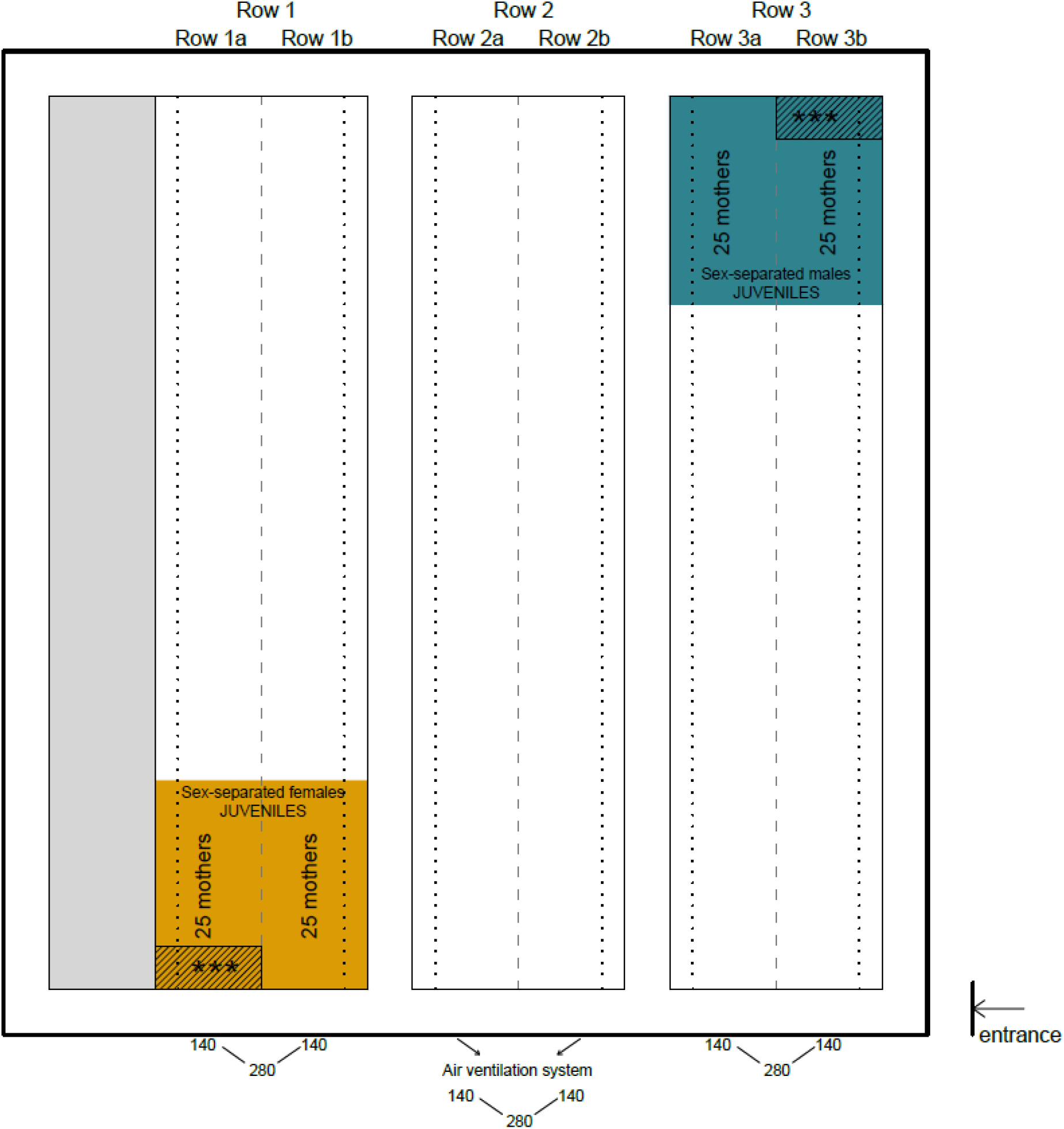
Maps of the farm and experimental design organization. **Scheme of room A (sex-separated animals).** The room has three double rows (1a, 1b, 2a, 2b, 3a, 3b) were mothers and their litters are located and an additional row (in grey) is used as a backup to replace animals of the main chain when needed. Each double row contains 280 mothers, each with their own litter (8 kits/litter). Litters are always re-arranged at day 0 to assure that all kits/mother have similar size. Air ventilation system works from the side towards the center of the room and in one direction only (see arrows in the scheme). Our experimental groups were of 50 mothers/experimental group and all eight kits/mother were either females or males for the sex-separated female and sex-separated male groups, respectively. The experimental groups (sex-separated female in orange and sex-separated male in green) were located according to the air ventilation system so that there is no air crossing. Weaning took place at day 30. Normal farm routing involves shifting of mothers to a different room to start the cycle again whereas the 8 kits / cage remain in the room until day 68, when they are sent to the slaughter house. In the experiment, mothers were taken out as usual farm routine, and the first **sampling point took place in room A at day 40,** in which 3 animals per group, each coming from a different mother, were used for the experiment. Animals were taken from the most distant sides of the room (indicated with ***). At day 40, 60 sex-separated males and 60 sex-separated females were shifted to rooms C and D respectively. The rest of the animals followed the normal farm routine.

**Scheme of room B (sex-combined animals).**
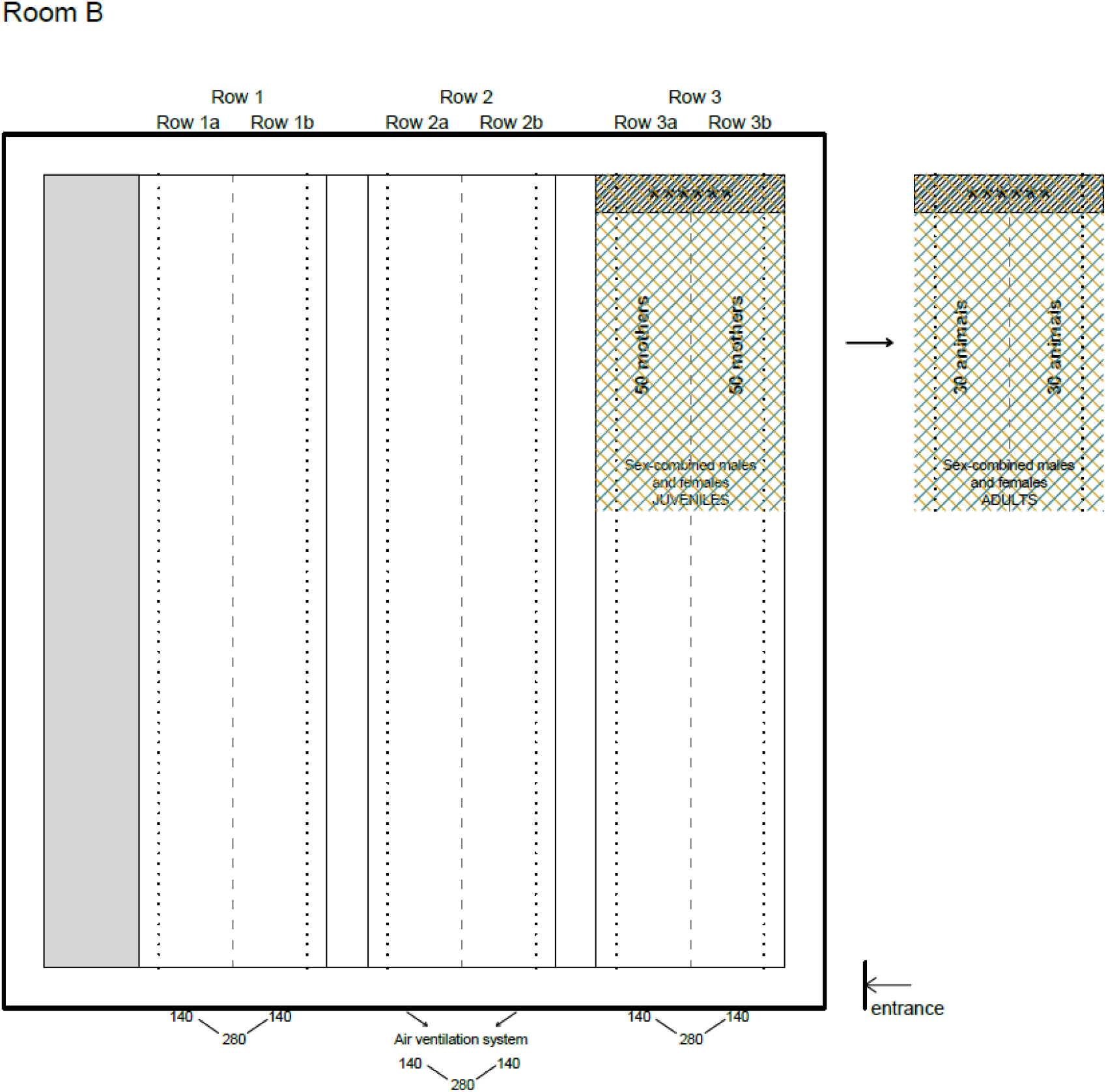
The room has the same features as room A. In this case, we employed 100 mothers, and each mother had 8 kits, 4 males and 4 females. Similarly to sex-separated condition, first sampling point took place at day 40, in which 6 animals (3 males and 3 females) (***), each coming from a different mother, were used for the experiment. Importantly, for the adults, animals were not shifted to another room; instead, they were kept in the same room to ensure common farm routine. Animals were re-arranged so that four individuals, two males and two females, were kept in each cage.

**Scheme of room C, D (sex-separated males and females respectively).**
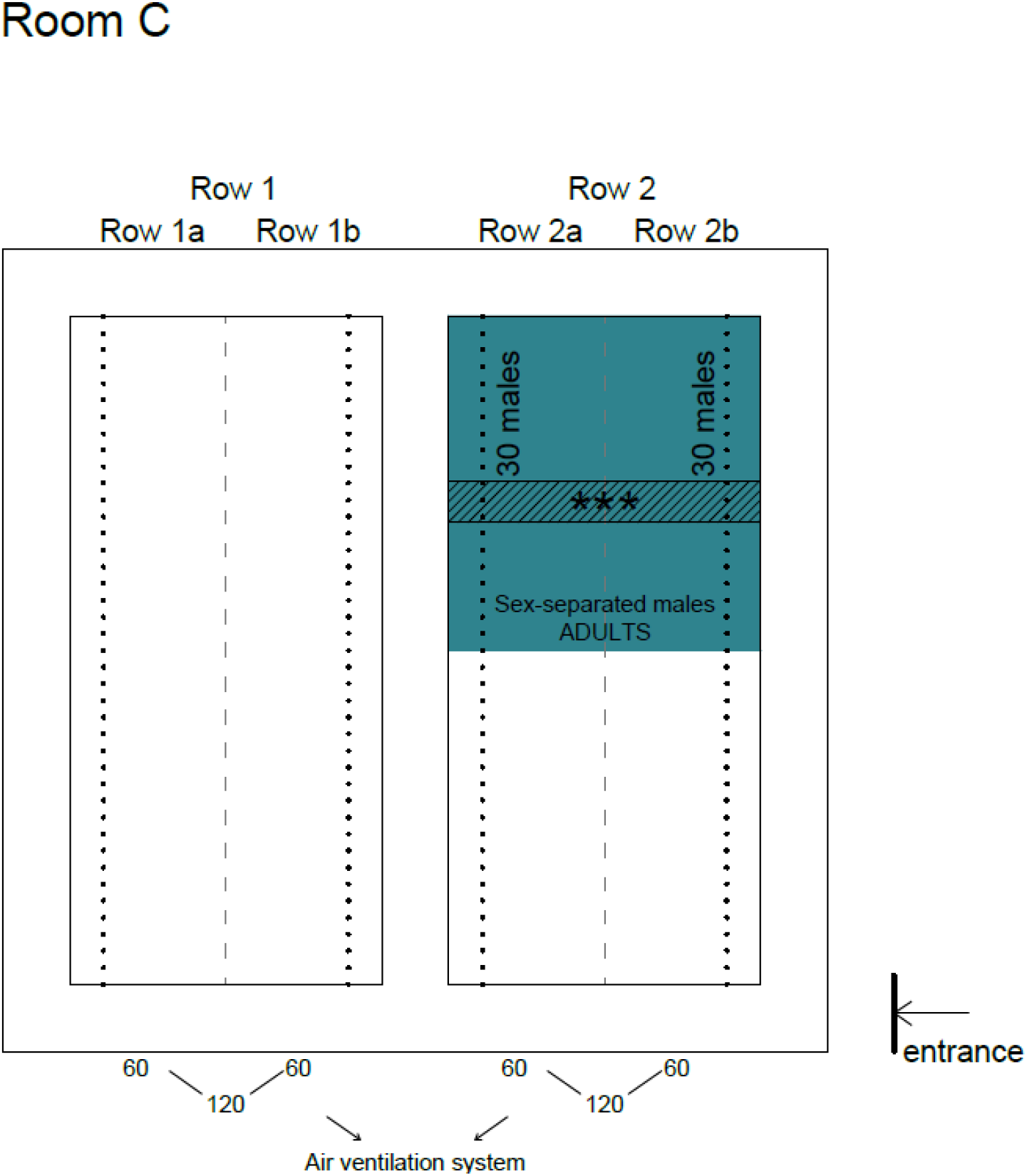
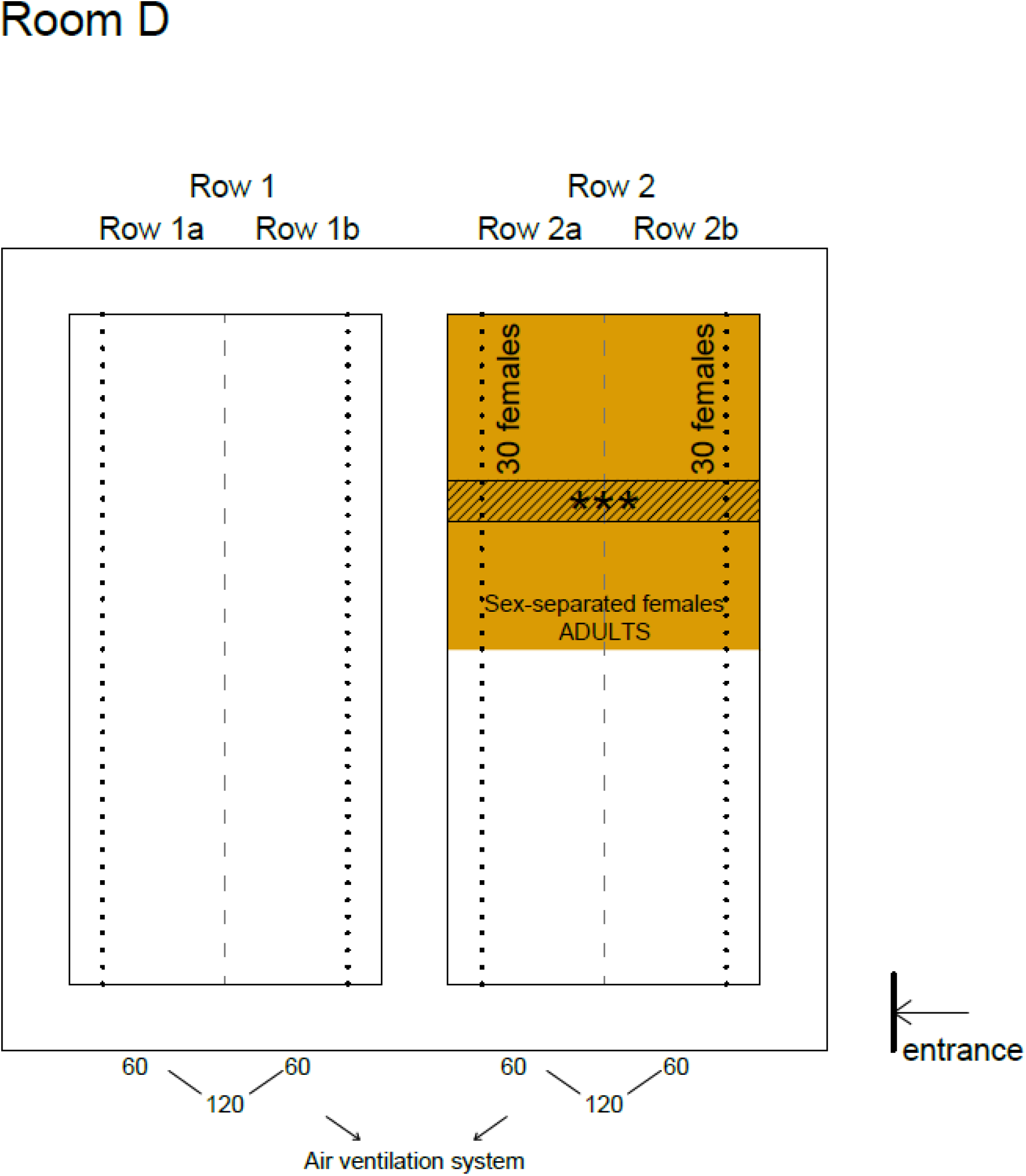
The room is considerably smaller than room A, containing three double rows (1a, 1b, 2a, 2b, 3a, 3b) and capacity for a total of 120 individuals. 60 sex-separated males and 60 sex-separated females were taken from room A at day 40 and placed in room C and room D, respectively. Four animals were placed / cage, assuring direct contact between individuals, and maintained until 6 months, in which three animals/experimental group from the most distant sides of the room (indicated with ***) were used for the experiment.

*Additional information:*

Some animals were eliminated due to health issues during the 6-month period. However, the outcome of the experiment was not affected thanks to the considerable high number of animals used to assure the ‘sex-separation’ and ‘sex-combined’ environments.

**Additional Figure 3.**
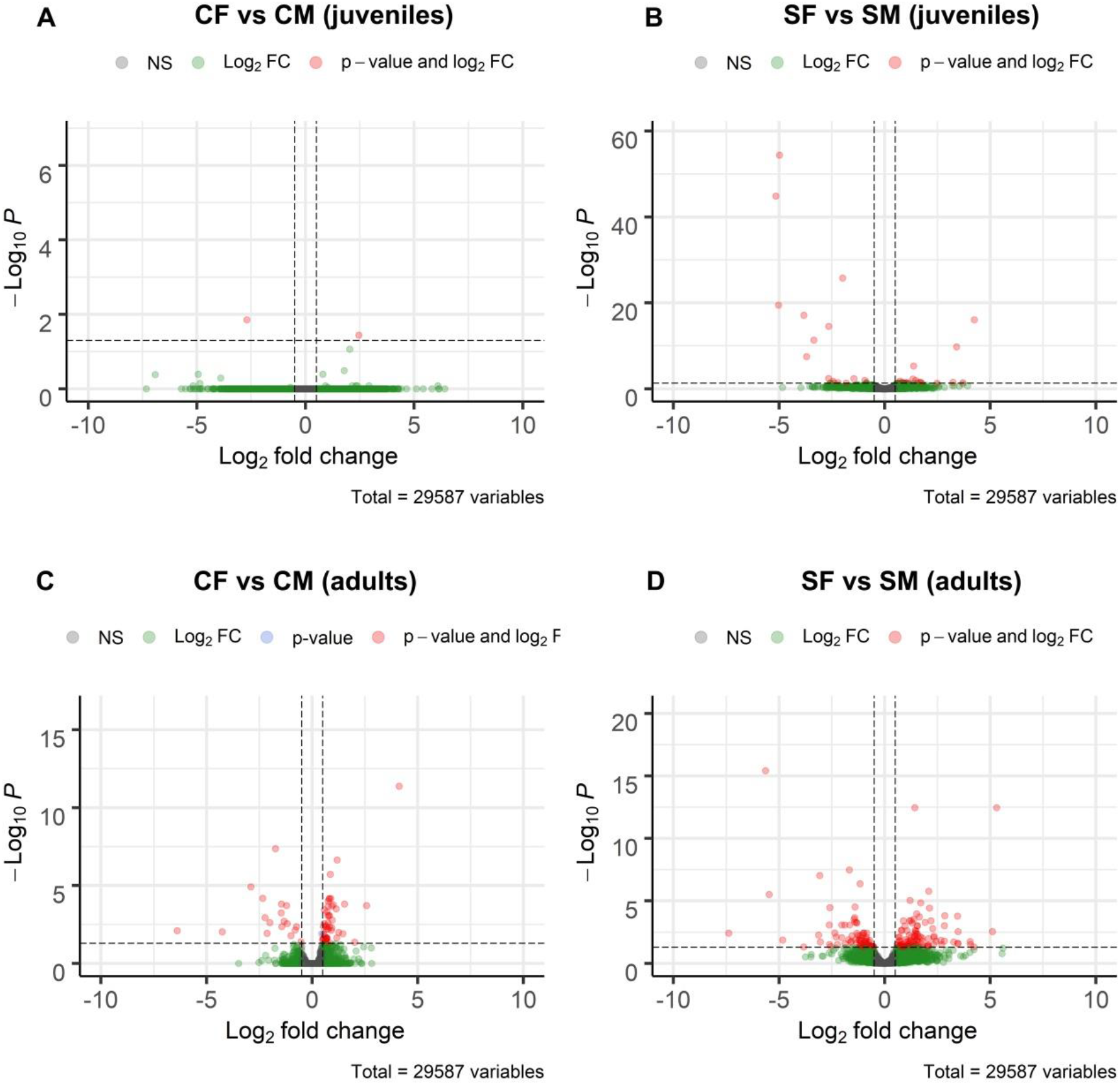
Volcano plots showing differentially expressed genes (fold change (FC), abscissa) and its significance (-log P-value, ordinates) between female and male VNO RNAseq samples: **(a)** juveniles sex-combined **(b)** juveniles sex-separated **(c)** adults sex-combined and **(d)** adults sex-separated. Genes are classified in four categories depending on their FC and FDR corrected p-value: i) grey = p-value > 0.01 and log2 FC between −0.5 and 0.5; ii) green = p-value > 0.01 and log2 FC < −0.5 or > 0.5; iii) blue = p-value < 0.01 and log2 FC between −0.5 and 0.5; and iv) red = p-value < 0.01 and log2 FC < −0.5 or > 0.5). CF: sex-combined females; CM: sex-combined males; SF: sex-separated females; SM: sex-separated males.

**Additional Figure 4.**
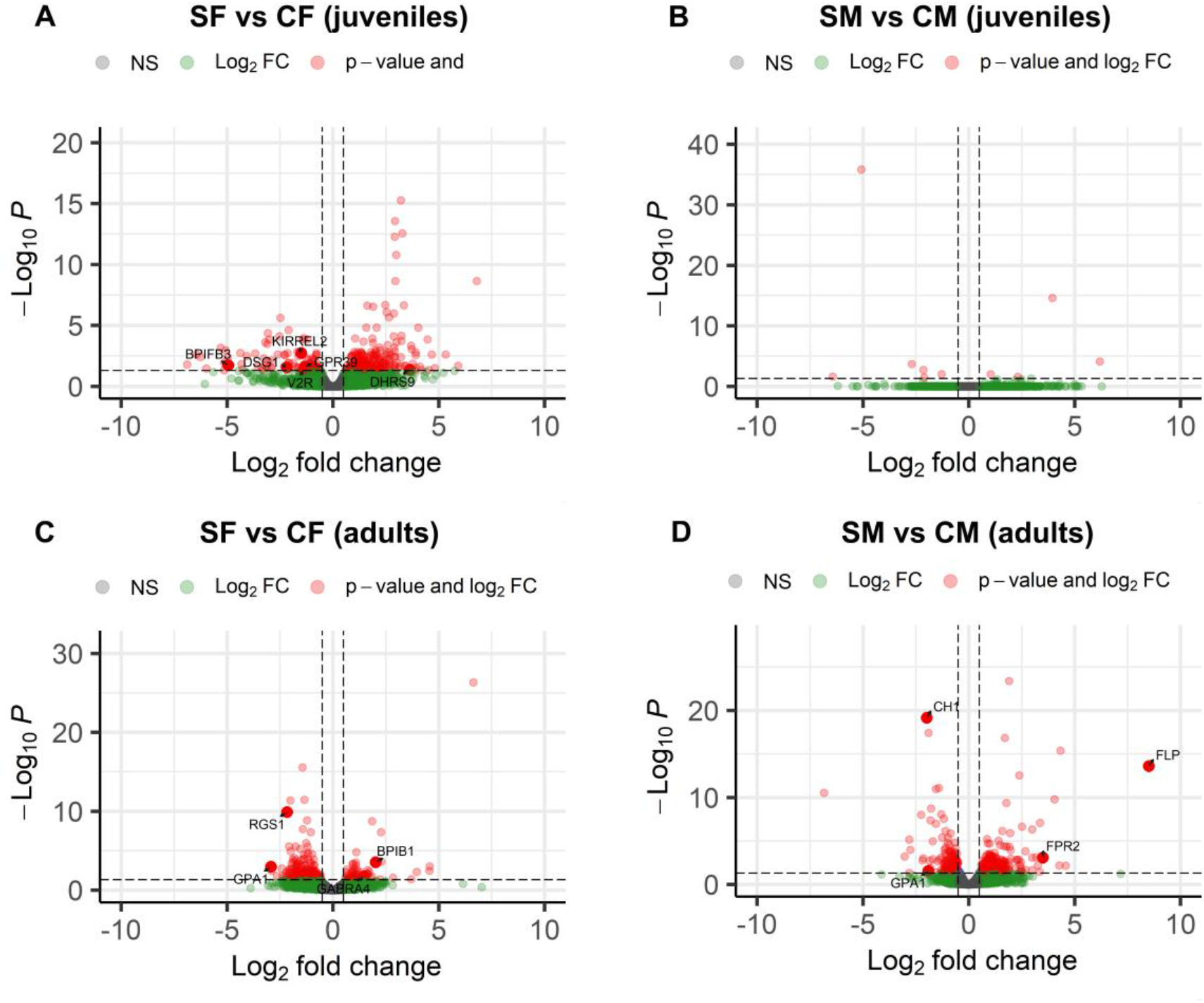
Volcano plots showing differential expression between sex-separated and sex-combined VNO RNAseq samples for the comparisons studied: **(b)** juvenile females **(c)** juvenile males **(d)** adult females **(e)** adult males. Each point in the plot represents a gene, with its log2 fold change (FC) in the x-axis and its log10 p-value in the y-axis. Genes are classified in four categories depending on their FC and FDR corrected p-value: i) grey = p-value > 0.01 and log2 FC between −0.5 and 0.5; ii) green = p-value > 0.01 and log2 FC < −0.5 or > 0.5; iii) blue = p-value < 0.01 and log2 FC between −0.5 and 0.5; and iv) red = p-value < 0.01 and log2 FC < −0.5 or > 0.5).

**Additional Figure 5.**
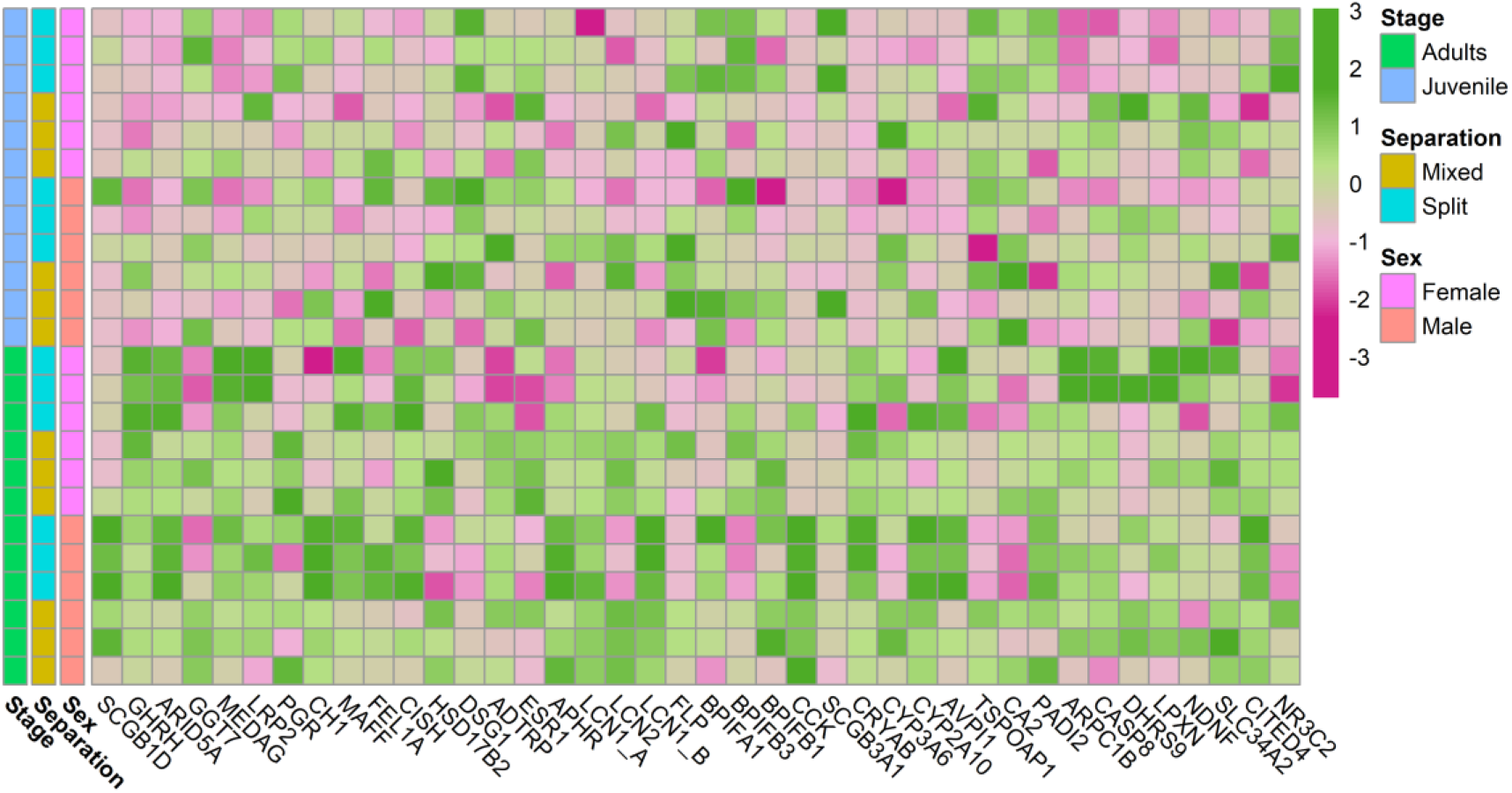
Heatmap of reproductive-related genes differencially expressed between comparisons, both in juveniles and adults

### ADDITIONAL TABLES

**Additional Table 1.**
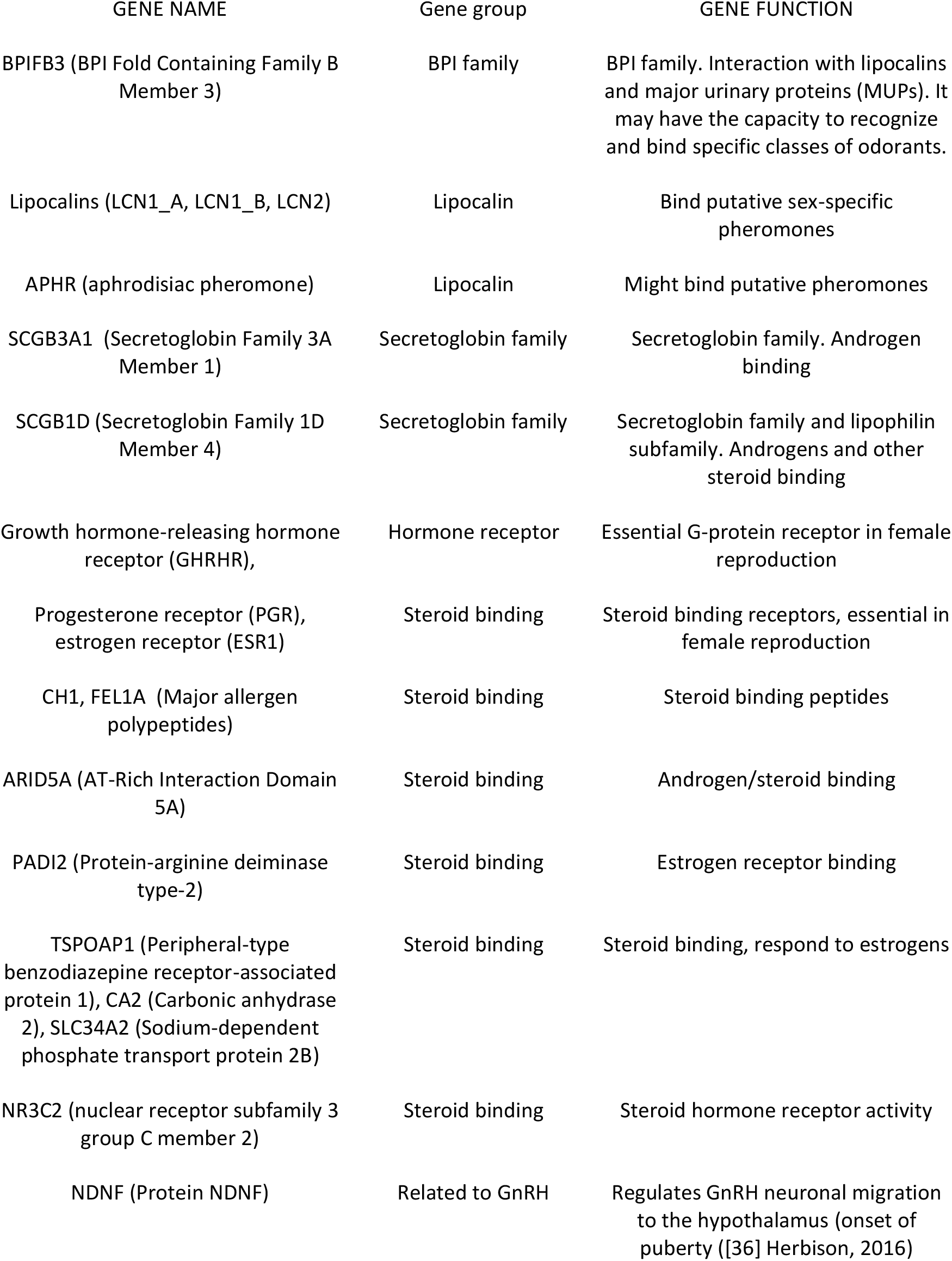
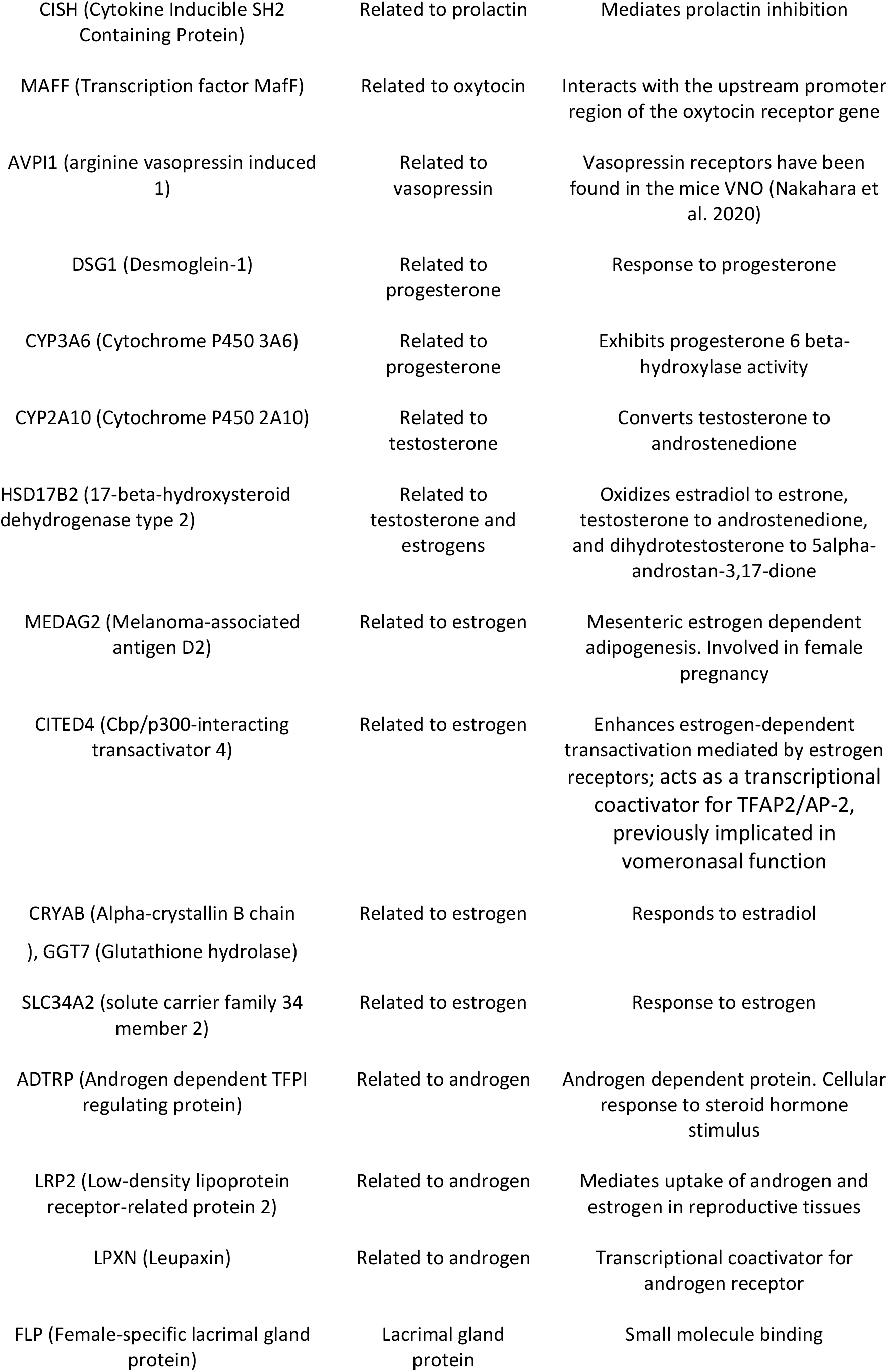
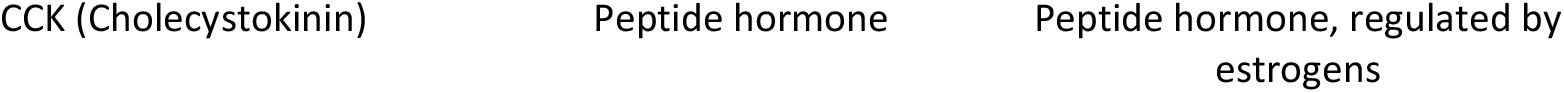
**Reproduction-related genes showing differential expression across socio-environmental conditions.**

### ADDITIONAL FILES

**Additional file 1:** List of differential expressed genes between males and females for both sex-separated and sex-combined animals, considering juveniles and adults.

**Additional file 2:** Overlapping genes considering DE female *vs* female

**Additional file 3:** List of differential expressed genes between sex-separated and sex-combined individuals, for both males and females, considering juveniles and adults.

**Additional file 4:** Overlapping genes considering DE sex-combined *vs* sex-separated

**Additional file 5:** List of differential expressed vomeronasal receptor genes between males and females for both sex-separated and sex-combined animals, considering juveniles and adults

**Additional file 6:** List of differential expressed vomeronasal receptor genes between sex-separated and sex-combined individuals, for both males and females, considering juveniles and adults.

**Additional file 7**: Enrichment analysis of the DEGs among experimental comparisons (SF *vs* SM, CF *vs* CM, SF *vs* CF, SM *vs* CM; both in adults and juveniles – only when the number of DEGs was sufficient to perform the analysis)

**Additional file 8:** List of genes belonging to the H2-Mv complex (vomeronasal receptors) in mice, and the corresponding genes found when blasted against the rabbit genome.

### Availability of data and materials

The RNA-seq raw reads analyzed in the current study are available on the National Center for Biotechnology Information (NCBI) Short Read Archive (SRA) database under accession number PRJNA720622. The datasets used and analyzed, as well as the analysis codes, are available from the corresponding author on reasonable request.

## Abbreviations

APHR: Aphrodisin
AQPs: Aquaporins
Bpifb3: BPI Fold Containing Family B Member 3
CCK: Cholecystokinin
CF: Combined female
CM: Combined male
DE: Differentially expressed
DEGs: Differentially expressed genes
ESR: Estrogen
FC: Fold change
FDR: False discovery rate
FEL1A: Major allergen I polypeptide chain 1
FLP: Female-specific lacrimal gland protein
FPRs: Formyl peptide receptors
GHRHR: Growth hormone-releasing hormone receptor
GO: Gene ontology
GPA1: Alpha subunit of the heterotrimeric guanine nucleotide-binding G protein
H2-Mv: A multigene family of non-classical class I major histocompatibility complex (MHC) genes
LCN1_B: Lipocalin 1B
LCN2: Lipocalin 2
MHC: Major histocompatibility complex
MSP: Male-specific submandibular salivary gland proteins
MUP: Major urinary protein
NCBI: National Center for Biotechnology Information
ORs: Olfactory receptors
PCA: Principal component analysis
PRG: Progesterone
RNAseq: RNA sequencing
SF: Separated female
SLC1A1: Excitatory amino acid transporter 3
SM: Separated male
SRA: Short Read Archive
V1R: Vomeronsal receptors type 1
V2R: Vomeronsal receptors type 2
VNO: Vomeronasal organ
VNS: Vomeronasal system
VR: Vomeronsal receptors
VSNs: Vomeronasal sensory neurons

## Acknowledgements

The authors thank COGAL SL (Pontevedra, Spain) for providing the facilities and animals employed in this study, as well as technical support—special thanks are due to Mª Carmen Diéguez for animal management support. We also gratefully thank Lucía Insua, Carlos Fernández and Andrés Blanco for technical and bioinformatic support and CESGA (Supercomputing Center of Galicia) for providing the necessary resources for the development of this work.

## Authors’ contributions

PM, PSQ and PRV conceived and designed the experiments and RNA-seq assay. PRV, JG, LQ and PSQ were involved in the experimental sampling. PRV and DR carried out the sequencing, transcriptomic analyses, bioinformatic work and data analysis. PRV wrote the original draft. DR and PM participated in the reviewing and critical analysis of the manuscript. All authors provided critical input and approved the final version.

## Corresponding author

Correspondence to Paula R Villamayor

## Ethics declarations

All experiments were approved by the ethical committee of the University of Santiago de Compostela, accordingly to the Directive of the European Union, 2010/63/EU, revising Directive 86/609/EEC, on the protection of animals used for scientific purposes.

## Consent for publication

Not applicable

